# SPHINGOLIPIDS AND Δ8-SPHINGOLIPID DESATURASE FROM THE PICOALGA *OSTREOCOCCUS TAURI* AND INVOLVEMENT IN TEMPERATURE ACCLIMATION

**DOI:** 10.1101/2023.05.16.541044

**Authors:** Toshiki Ishikawa, Frédéric Domergue, Alberto Amato, Florence Corellou

## Abstract

Sphingolipids are crucial components of cell membranes. Sphingolipid Δ8-unsaturation is more specific to plants and is involved in the regulation of stress responses. The structure and functions of sphingolipids in microalgae are still poorly understood. *Ostreococus tauri* is a minimal microalga at the base of the green lineage, and is therefore a key organism for understanding lipid evolution. The present work reports the characterisation as well as the temperature regulation of sphingolipids and Δ8-sphingolipid desaturase from *O. tauri*. Complex sphingolipids are glycosylceramides with unique glycosyl moieties encompassing hexuronic acid residues, reminiscent of bacterial glucuronosylceramides, with up to three additional hexose residues. In contrast, the ceramide backbones show limited variety, with dihydroxylated C18/C18:1^EΔ8^ sphingoid bases and C16:0 fatty-acyl chain being the main compounds.

The sphingolipid Δ8-desaturase from *O. tauri*, although phylogenetically related to plant homologues has a substrate preference similar to the diatom homologue. Both sphingolipid Δ8-desaturase transcripts and sphingolipid Δ8-unsaturation are regulated in a temperature-dependent manner being higher at 14°C than 24°C. Overexpressing the sphingolipid Δ8- desaturase in *O. tauri* at 24°C results in higher sphingolipid unsaturation and impairs the increase in cell size, structure and chlorophyll. In particular, the cell-size defect is not detected in cells acclimated to 14°C and is furthermore suppressed upon transfer from 24°C to 14°C. Our work provides the first functional evidence for the involvement of sphingolipid Δ8-unsaturation for temperature acclimation in microalgae, suggesting that this function is an ancestral feature in the green lineage.

## INTRODUCTION

Sphingolipids (SLs) are ubiquitous lipids occurring in all eukaryotes as well as in a few bacteria (Hannich et al., 2011). SLs structure membranes and, along with sterols and glycosyl phosphatidylinositol (GPI) anchored proteins, are essential for the dynamics of membrane microdomains (Lingwood and Simons, 2010). SLs have been shown to be importantly involved in biotic and abiotic stresses including temperature responses (Fabri et al., 2020; Huby et al., 2020; Wang et al., 2021). Moderately soluble SLs metabolites such as long-chain base (LCB) and ceramides (Cer) play critical roles as ligands that bind to, and regulate the activity of, enzymes and signaling proteins such as kinases and membrane receptors (Breslow and Weissman, 2010). Moreover, the balance between non-phosphorylated and phosphorylated forms of LCB and Cer has been shown to be involved in programmed cell death in animals, plants and yeasts (Ali et al., 2018).

SLs encompass a ceramide backbone consisting of a long-chain sphingoid base (LCB), commonly a C18 acyl-chain with hydroxyl groups at C1 and C3, to which a fatty acid (FA) chain of various length is amidified (at C2) (Sperling and Heinz, 2003). Ceramide diversity arises from changes in chain length, methylation, hydroxylation and/or degree of desaturation of both the LCB and FA moieties. Mono- or pluri-hexoses in glycosylceramides (GlyCers), phosphoryl in ceramide-phosphates or inositol-phosphate group in phosphoinositol ceramides (IPCs) further substitutes the primary hydroxyl of the sphingoid base. Monohexosylceramides are the simplest glycoSLs. GlyCers occur widely including in animals while IPCs are common to plants, fungi, and some protists (Hsu et al., 2007; Cacas et al., 2013; Vítová et al., 2022; Yamashita et al., 2022). GlycosylIPC (GIPC) result from additional transfer of saccharide residues to phosphoinositol ceramides (IPCs) by glycosyltransferases. Plants complex sphingolipids are glucosylceramides consisting of only one glucose (GlcCers), and GIPCs consisting of highly diverse and complex headgroups defining eight series (Gronnier et al., 2016). Noteworthy, plant GIPCs encompass glucuronic acid (GlcA), which is added to the IPC backbone by inosiltolphosphate ceramide glucuronosyltransferase (Cacas et al., 2012; Rennie et al., 2014).

Overall, the core metabolic pathway of SLs is well conserved in eukaryotes (Mashima et al., 2019). SL *de novo* synthesis starts with the condensation of serine and palmitoyl-CoA, catalysed by the serine palmitoyltransferase (SPT) in the ER, and produces 3- ketosphinganine, which has a hydroxyl at C1, an amine group at C2 and a ketone at C3. Further reduction by the 3-ketosphinganine reductase (KSR) generates sphinganine (d18:0, dihydrosphingosine), which is next N-acylated by the ceramide synthase (CS) and yields ceramides. Headgroup attachment takes places in the Golgi. Concerning LCB modifications, C9-methylation appears specific to fungi while LCB hydroxylation and desaturations are widespread. C4 hydroxylation and Δ4-desaturation of sphinganine are mutually exclusive, as the corresponding enzymes in charge of these modifications, namely C4-hydroxylase and SL Δ4-desaturase (SLD4), respectively, are competing for the same substrate. The resulting 4- hydroxysphinganine (t18:0, phytosphingosine) and 4-sphingenine (18:1^4E^, sphingosine) are commonly encountered in eukaryotes. LCB Δ8-unsaturations has been reported from plants, algae, moss and fungi and may be concomitant with C4-modifications yielding either t18:1^E8^ or d18:2 ^E4,E/Z8^ (Stonik and Stonik, 2018; Mashima et al., 2019; Liu et al., 2021; Haslam and Feussner, 2022).

Sphingolipid Δ8-desaturases (SLD8) have first been characterised from plants (Sperling et al., 1998). SLD8 encompass N-terminal (Nt) fused cytochrome b5 domain (CytB5). SLD8 from fungi generates Δ8-*E* isomers while plant SLD8 generate both *Z* and *E* Δ8 isomers with different efficiencies depending on the species (Li et al., 2016). The ratio between *Z*- and *E-*Δ8 isomers was proposed to be related to chilling resistance and, to some extent, to aluminium tolerance (Imai et al., 1997; Li et al., 2016). For microalgae, the only SLD8 functionally characterised to date is TpDesB from the diatom *Thalassiosira pseudonana* (Tonon et al., 2005). TpDesB introduces a Δ8 in the *E* configuration and displays a strong preference for dihydroxylated LCB substrates.

Knowledge about microalgal SLs is poor especially in view of the phylogenetic diversity of these organisms. GlyCers were detected in a large number of microalga species (Arakaki et al., 2013; Li et al., 2017; Stonik and Stonik, 2018). The first structural study was carried out on the green microalgae *Tetraselmis sp*. and unveiled original GlcCer in which a plant-related LCB (d18:2^4E,8E^) is combined with a FA bearing modification only reported from fungi (2- hydroxy-Δ3-unsaturated FA) (Arakaki et al., 2013). Recently, structural analyses of SLs in major species of microalgae from the Stramenopiles, Alveolates, Rhizaria supergroup (SAR) have been achieved (reviewed in (Stonik and Stonik, 2018)). The results highlighted unique features of microalgal GlyCers such as diverse glycosyl moieties consisting of di- to tri- saccharide residues (diatoms) or original monoresidue such as galactose and sialic acid (haptophytes) as well as fungal-like C9 methylation and/or tri to tetra unsaturation of LCBs.

*Ostreococcus tauri* is a marine picoalga that emerged in the green lineage 1.5 billion years ago. *O. tauri* corresponds to the smallest photosynthetic eukaryote (< 2µm) and defines with other related picoeukaryotes the Mamiellophyceae class and Mamiellales order (Courties et al., 1994; Marin and Melkonian, 2010). Among Mamiellophyceae, *O. tauri* has the most reduced genome (13 Mb) and the simplest cellular organization (Derelle et al., 2006; Henderson et al., 2007). In the utra-compact cell, the unique chloroplast occupies about 50% of the cell volume and is in contact with most of other organelles. *O. tauri* glycerolipidome has been characterised in details unveiling features reminiscent of both the SAR and Archaeplastida supergroups (Degraeve-Guilbault et al., 2017). In particular, *O. tauri* and related species produces very-long chain polyunsaturated FA (VLC-PUFAs, C>20) that are hallmark of SAR microalgae. The Mamiellophyceae FA desaturases crucial for VLC-PUFA synthesis, namely Δ6-, Δ5-, Δ4-desturases (Des), have been functionally characterized (Domergue et al., 2005; Tavares et al., 2011; Vaezi et al., 2013; Degraeve-Guilbault et al., 2020). In contrast to other ancient eukaryotes which encompass ER located Δ6-desturases using lipids as substrate, Mamiellophyceae display animal-like acyl-CoA Δ6-Des that uses acyl-CoA as substrate (ER located) as well as plastidic FA Δ6-Des that are plastid located and uses lipid substrates (Domergue et al., 2005; Degraeve-Guilbault et al., 2020).

In the present work, we aimed at gaining insight into the sphingolipid structure/function in *O. tauri*. We therefore functionally characterised the SLD8 and SLs from *O. tauri* and further provide evidences for their involvement in temperature acclimation.

## RESULTS

### Phylogenetic analysis of sphingolipid **Δ**8 desaturase candidates

Among CytB5-fused desaturase we previously retrieved, the accession XP_022841141 had been identified as a putative SLD8 (Degraeve-Guilbault et al., 2020). Bayesian phylogenetic analysis using amino acid (AA) sequences of all CytB5-fused desaturases from Mamiellales shows that the putative *O. tauri* SLD8 AA sequence robustly clusters (posterior probability PP = 1.00) with other Mamiellales putative SLD8 and is related (posterior probability PP = 0.75) to the clade of Mamiellales plastidic Δ6-Des (Fig. 1) (Degraeve-Guilbault et al., 2020). The acyl-CoA Δ6-Des clade robustly (1.00) clusters basally to the clade containing both the SLD8 and the plastidic Δ6-Des. The phylogenetic analysis run with SLD8 from other organisms shows that the Mamiellales clade, together with its sister clade (PP= 0.70) containing Charophyte sequences, sits basally (PP = 1.00) to a robust clade (PP = 1.00) containing characterised SLD8 from Bryophytes and higher plants (Fig. 2). The *Nymphaea thermarum* sequence defines the monocot/eudicot separation, as expected following the Paleoherb hypothesis (Doyle and Donoghue, 1993; Igersheim and Endress, 1998). Using a NJ phylogenetic inference, Li and collaborators reported a clear dichotomy between plant SLD8 preferentially producing *Z* and *E* isomers (Li et al., 2016). From our analysis this dichotomy was not as clear, though *Z*>*E* eudicots robustly (P = 1.00) cluster together (with one exception, *Prunus persica*), whereas the *E*>*Z* are more scattered in the tree, still showing a pattern. Considering sequences from the distant microalgal lineage SAR, sequences cluster together, with the exception of the two diatoms species *Nitzschia inconspicua* and *Thalassiosira pseudonana*. The only cryptophyte in the analysis (*Guillardia theta*), has possibly destabilised the analysis (PP = 0.55). *T. pseudonana* SLD8 is the only SAR sequence that has been functionally characterised to date and was shown to exclusively produce *E* isomer (Tonon et al., 2005).

**Figure 1.**
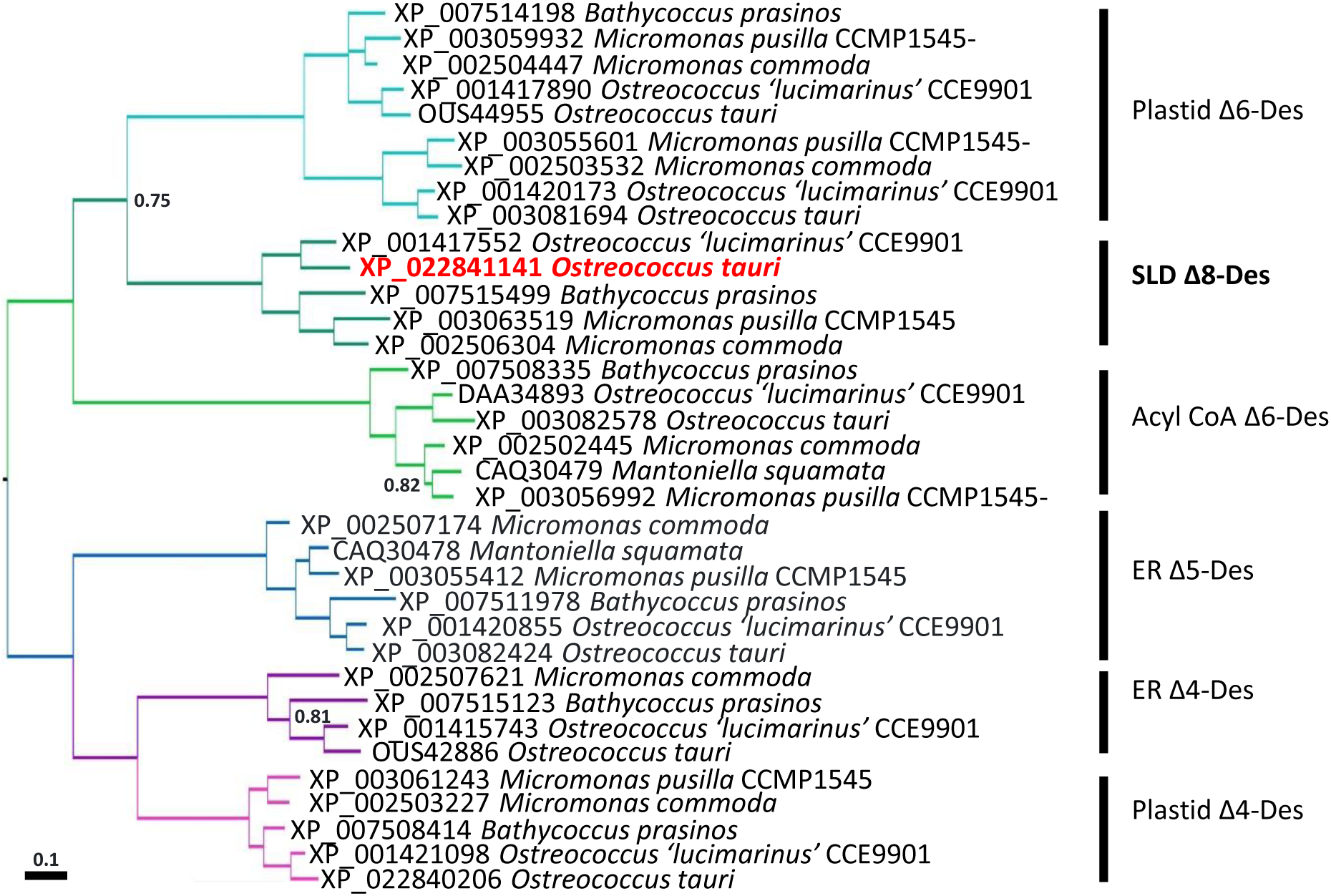
Bayesian phylogenetic tree of Mamiellales desaturases. The tree was drawn to scale in substitutions per site (scale bar on the figure). Posterior Probabilities (2.000.000 generations) greater than 0.95 are not reported, the others are indicated by the nodes. Vertical bars indicate protein function. The *O. tauri* accession is highlighted in red

**Figure 2.**
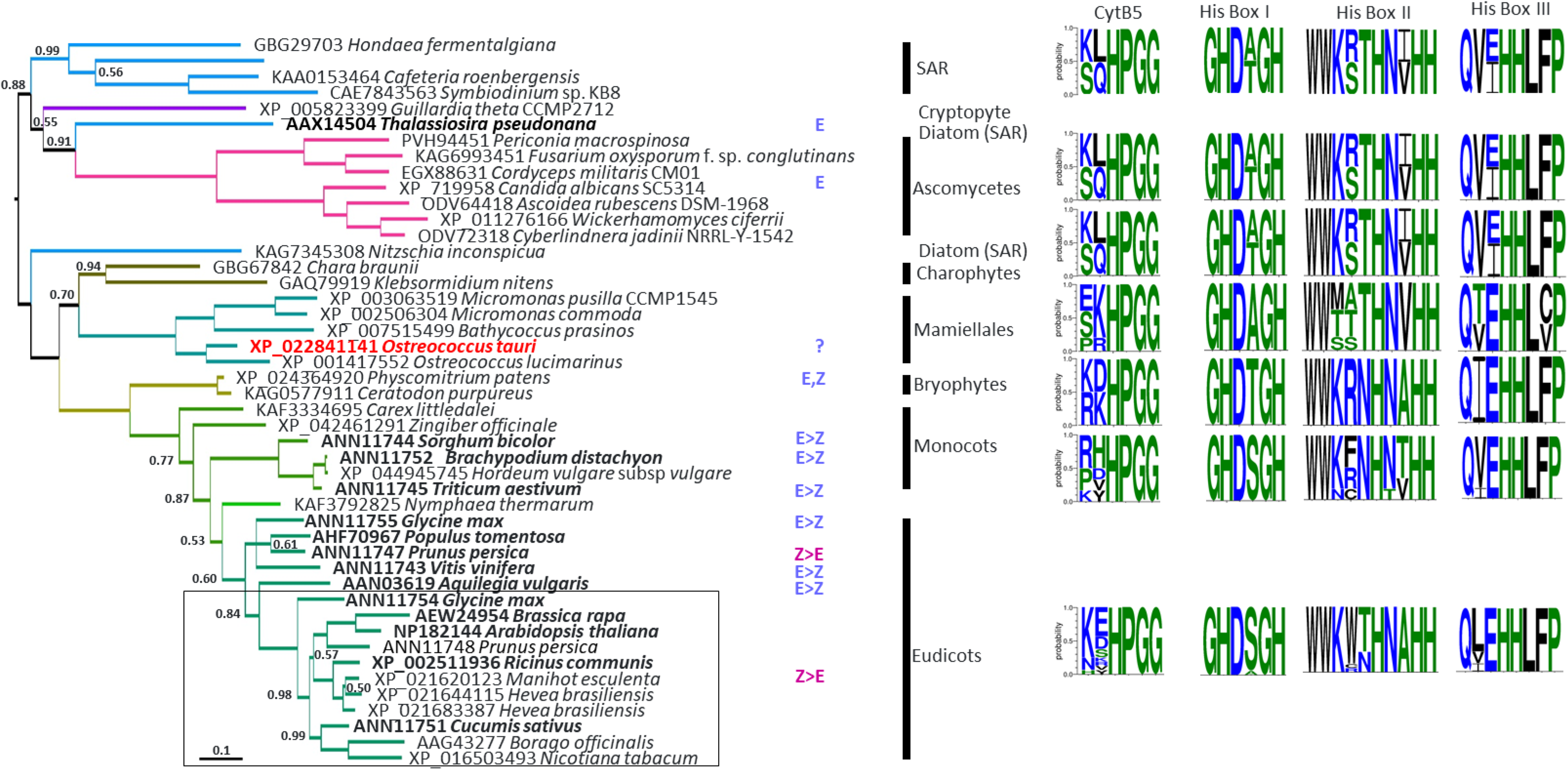
Bayesian phylogenetic tree of sphingolipid Δ8 desaturases. The tree was drawn to scale in substitutions per site (scale bar on the figure). Posterior Probabilities (2.000.000 generations) lower than 1.00 are indicated by the nodes. Vertical bars indicate taxonomic assignment, also visualised by brench colors. The preference of the enzyme to catalyse the *E*- or *Z*-isomer is indicated on the figure and is based on functional characterisation of the protein (Tonon et al 2005, Oura et al 2008, Li et al 2016). According to Steinberg (2021) the occurrence of *E* and *Z* in *P. patens* is indicated. Sequence logos of the previously identified histidine boxes are presented on the right-hand side. The letter hights are drawn in scale to the probability of occurrence in the sequences used for the sequence logo production (SAR n = 4; Ascomycetes n = 7; Charophytes n = 2; Mamiellales n = 5; Bryophytes n = 2; Monocots n = 7; eudicots n = 16).

### Subcellular localization of SLD8 candidates

In order to possibly up-date the start codon provided in the NCBI/JGI accessions we searched for additional possible start codons 5’upstream of, and in frame with, the annotated sequence (Fig. S1). Putative start codons occurred for *O. tauri*, *Ostreococcus* sp. RCC809 and *Micromonas pusilla*. For *O. tauri*, we successfully amplified the corresponding sequence from cDNA. From subcellular localisation prediction (PredAlgo) fair to high scores for chloroplastidic target peptide (cTP) were obtained for all Mamiellales sequences exception made of *Bathycoccus prasinos*. In particular, the 33 AA extended OtSLD8 sequence displayed a cTP score of 3.11 over 5 (Fig. S1). We therefore considered a short version and a long version of SLD8 for subsequent analyses (short-SLD8 and long-SLD8 thereafter). Since sphingolipid desaturases are commonly assumed to be ER located, we wanted to assess the subcellular localisation of both short-SLD8 and long-SLD8. To this end *Nicotiana benthamiana* was used as heterologous host for the expression of Cterminal-YFP-fused-SLD8 (Degraeve-Guilbault et al., 2020; Degraeve-Guilbault et al., 2021). The short-SLD8 was unambiguously ER located while the long-SLD8 localised at chloroplasts (Fig. 3).

**Figure 3.**
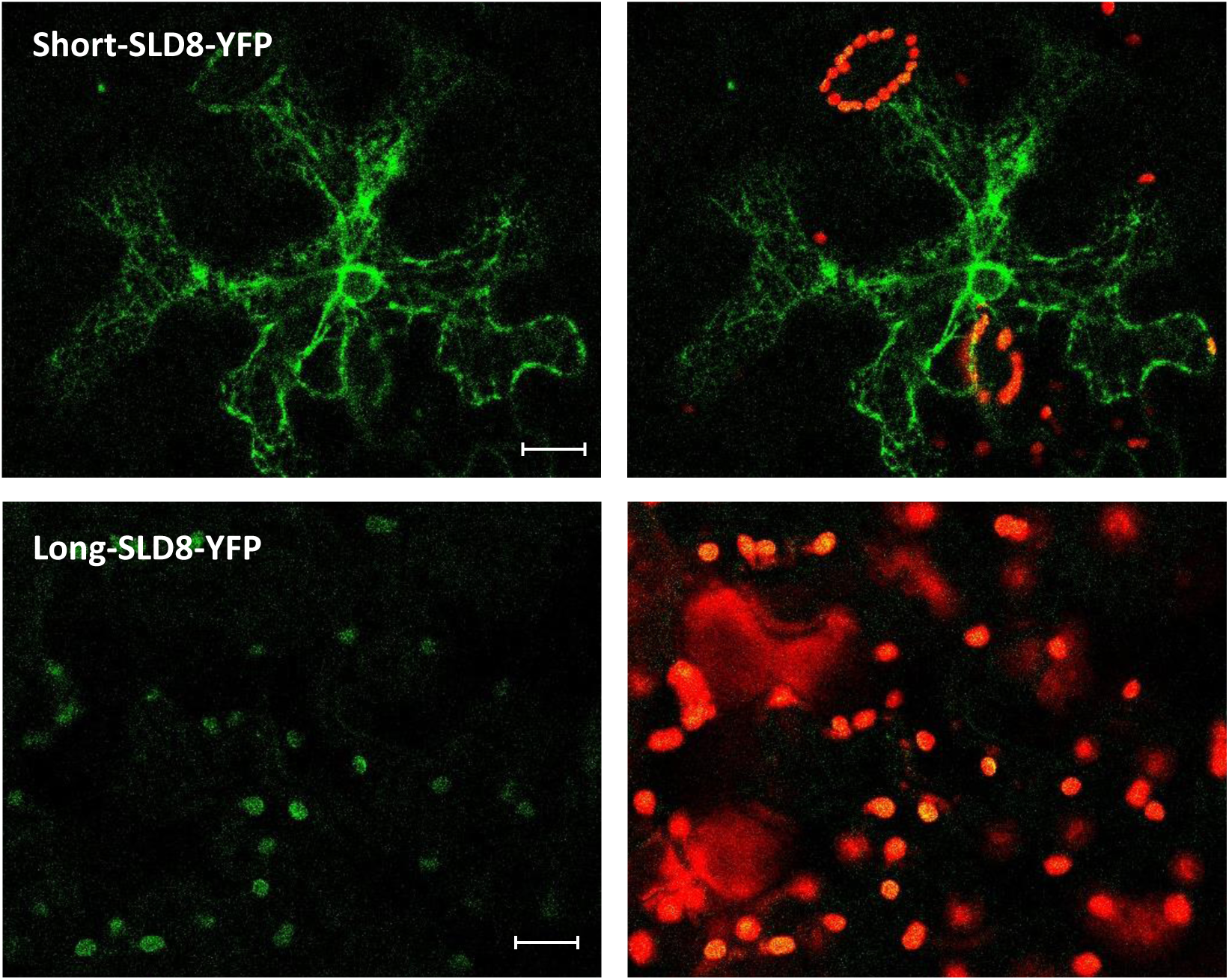
Sub-cellular localisation of *O. tauri* sphingolipid Δ8-desaturase short and long versions in *Nicotiana benthamiana*. Confocal images of *Nicotiana benthamiana* leaves transiently transformed with OtSLD8-YFP constructs. *O. tauri* sphingolipid Δ8-desaturase short and long versions were cloned into the pK7YWG2 destination vector, the resulting constructs transferred into the *Agrobacterium tumefaciens* strain GV3101, and transformants used for transient expression in leaves. Images were taken 2 days post infiltration. Left, YFP signal; Right, merged of YFP signal and chlorophyll autofluorescence. Bars = 20µm.

### SLD8 enzymatic activity

#### O. tauri LCB and ceramides

LCB and ceramides analyses of *O. tauri* unveiled that sphinganine and its corresponding *E*-Δ8-desaturation product sphingenine were the main LCB, and that they are associated with the FA C16:0 in ceramides. (Fig. 4, Table S1). These results suggested that the putative OtSLD8 possibly displayed a preference for sphinganine and *E*-isomers.

**Figure 4.**
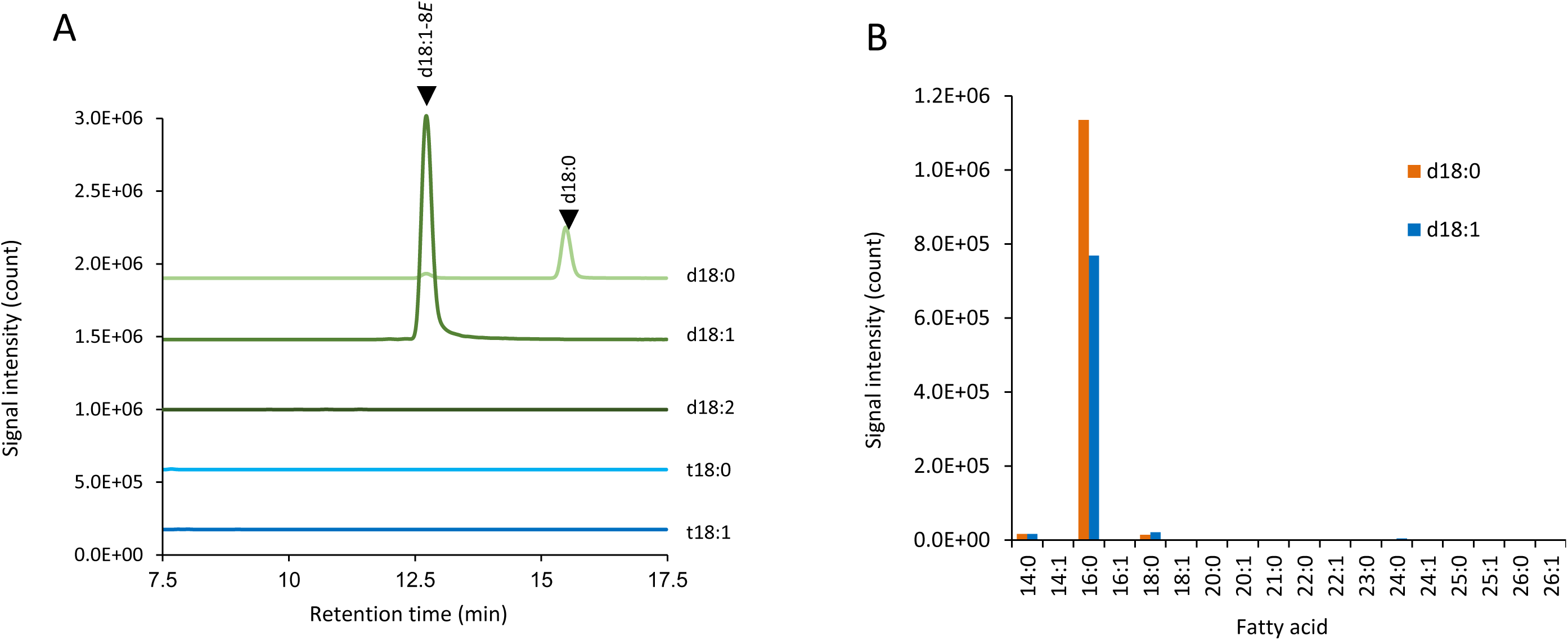
Profile of total LCB and fatty acyl moiety of free ceramides in wild-type *O*. *tauri*. **A**. LC-MS/MS profile of total LCB liberated from whole cells of *O*. *tauri*. LCBs were analysed as the NBD-derivatives by LC-MS/MS and the MRM chromatograms of the major 5 species conservatively observed in plants are shown at the same scale of absolute intensities of MS signal intensities. **B**. Fatty acid profile of free ceramides. Free ceramides composed of d18:0 or d18:1 LCB and one of various fatty acids were detected by LC-MS/MS by the targeted MRM mode. The MRM transitions are shown in Table S1.

#### Sphingolipid desaturase activity characterisation

Heterologous expression of OtSLD8 in yeast was carried out in order to validate the sphingolipid desaturase activity (Table 1). Both short- and long- *SDL8* were expressed in different *Saccharomyces cerevisiae* strains available from previous studies, along with SLD1 from *Arabidopsis thaliana* as positive control and an empty vector as negative control. *S. cerevisiae* strains allowed to test specific substrate: the wild-type strain provides t18:0, the *sur2Δ* mutant d18:0 and the *sur2Δ* expressing the *LCB Δ4 desaturase* gene of *Komagataella pastoris* provides d18:1^Δ4E^ (Haak et al., 1997; Tonon et al., 2005). Further note that the *pGAL1* inducible system was used in addition to the constitutive expression system (promoter *PGK1)* as expression of *long-SLD8* slowed-down the growth of yeasts. Expression of both the long and short versions of OtSLD8 resulted in five times more abundant unsaturation products from sphinganine than from 4-hydroxysphinganine, reflecting a preference of SLD8 for dihydroxylated substrates over trihydroxylated substrates. (Table 1) (Fig. S2). Considering the inducible expression, this activity was the higher for the short- OtSLD8. This contrasted with AtSLD1, which showed comparable unsaturation efficiency for sphinganine and 4-hydroxysphinganine. Considering the constitutive expression of the *short- OtSLD8*, both saturated and monounsaturated dihydroxylated substrates (d18:0, d18:1^Δ4E^) appeared equally well accepted while SLD1 has a clear preference for d18:1^Δ4E^ (sphingosine). Most interestingly, only *E* stereoisomers of the Δ8-double bond were produced by both the long- and short-OtSLD8, whereas *E* and *Z* isomers were generated by AtSLD1 (Fig S2).

**Table 1.**
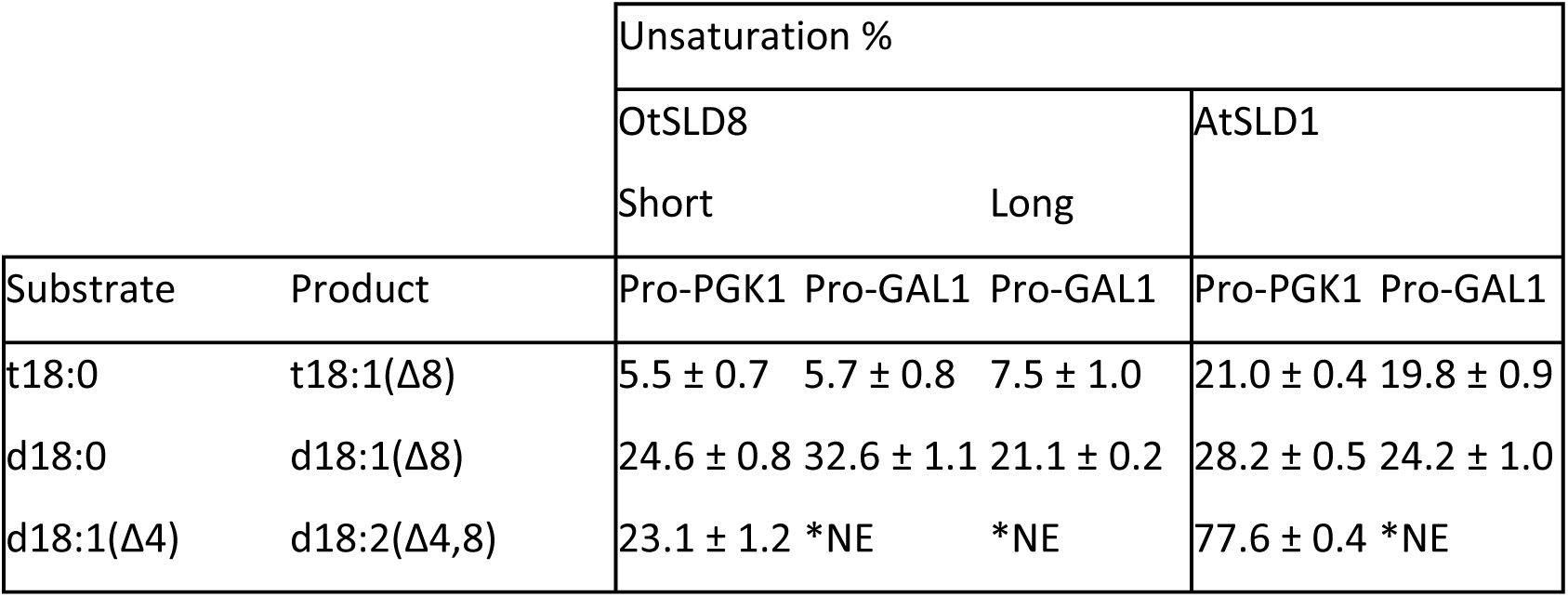
OtSLD8 LCB desaturase activity in S. cerevisiae.

#### Search for additional desaturase activity

*O. tauri* has the capacity of producing a peculiar fatty acid 18:5^Δ3,6,9,12,15^ which is commonly found in species from the SAR supergroup (Degraeve-Guilbault et al., 2017). C18:5 is recognized as a hallmark of galactolipids (plastidic lipids) and is assumed to be derived from the activity of a previously unidentified Δ3 desaturase using 18:4^,6,9,12,15^ as substrate. We recently characterised the first plastidic Δ6-Des from *O. tauri* and showed that their expression in *N. benthamiana* resulted in the accumulation of the Δ6-desaturation products 18:3n-6 18:4n-3 especially in plastidic lipids (Degraeve-Guilbault et al., 2020). As heterodimerization of closely related desaturases and desaturase bifunctionality have both been reported, we wanted to investigate whether SDL8 could display an additional activity when co-expressed with Δ6-Des in *N. benthamiana* (Sperling et al., 2003; Lou et al., 2014). We therefore overexpressed *long-SLD8* together with each of the *pΔ6-Des (pΔ6-Des1 and pΔ6-Des2*, previously referred to as Ot05 and Ot10 respectively) (Fig. 5A) and *Short-SLD8* with the ER *acyl-CoA-Δ6-Des* (previously referred to as Ot13) (Fig. 5B) (Degraeve-Guilbault et al., 2020). Consistently with previous reports, the transient overexpression of *Δ6-Des* in *N. benthamiana* resulted in the production of 18:3n6 and 18:4n3 from 18:2n6 and 18:3n3 respectively with the greatest conversion observed for pΔ6-Des1 (Fig. 5A). However, expression of OtSDL8 alone or in combination with other OtΔ6-Des did not result in other changes in the fatty acid profiles of *N. benthamiana*.

**Figure 5.**
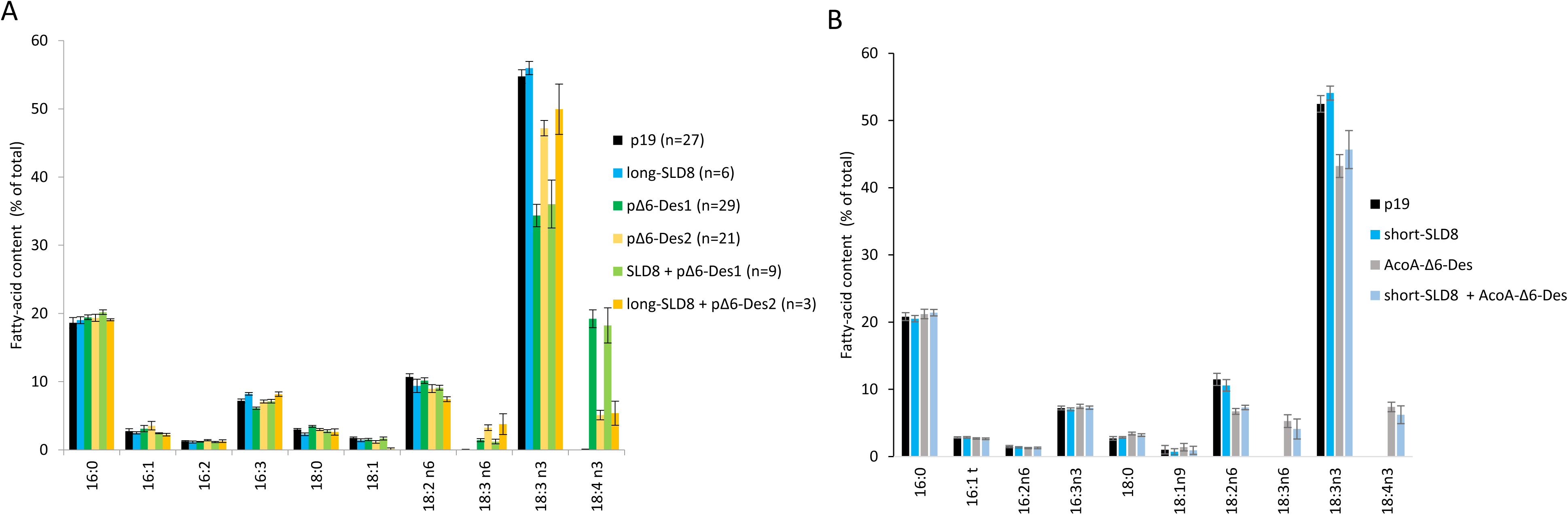
Fatty-acid profile of *N. benthamiana* expressing SDL8 alone and together with Δ6-desaturases. The anti-silencing protein p19 was used in all infiltrations. **A**. The long-SLD8 (plastid located) was expressed alone and together with the plastidic Δ6 desaturases pΔ6-Des1 and pΔ6-Des2. The number of replicate is indicated. **B**. The short version of SLD8 (ER located) was expressed alone and together with the ER Acyl-CoA-Δ6-Des (ACoA-Δ6-Des) (n=3). Means and standard errors are shown.

Altogether, our results show that SLD8 activity is specific to SLs and introduces a Δ8 desaturation exclusively in the *trans*-configuration into sphinganine, yielding 8E-sphingenine (d18:1^E8^).

### SLD8 functional analyses in *O. tauri*

SLs and SLD8 have been reported to be involved in temperature acclimation across eukaryotes and in particular in plants (Chen et al., 2012; Zhou et al., 2016; Fabri et al., 2020). We therefore chose to investigate the relationship between SL-Δ8-unsaturation and temperature acclimation in *O. tauri.* SLD8 overexpressing lines (SLD8 OEs) were generated to gain insight into the physiological relevance of SL-Δ8-unsaturation (Fig. S4). Cells synchronized by 16h light/8h dark cycles were acclimated for at least 3 sub-culturing rounds to high (24°C) and low temperature (14°C). Warming and cooling were carried out 3 to 4 hours after the light on by transferring acclimated culture from 14°C to 24°C and conversely. These conditions have been previously identified for allowing oscillatory expression of desaturases (peak in the morning), as well as studying temperature-dependent changes of glycerolipids and of desaturase expression (Monnier et al., 2010; Degraeve-Guilbault et al., 2021; Kay et al., 2021) (Fig. S4).

#### Effect of temperature on SLD8 expression and LCB unsaturation

SLD8 transcript abundance was monitored at 24°C and 14°C (acclimated cells) and 0.5 and 4 hours after a temperature shift that was achieved within 3 minutes (see methods) (Fig. 6). The transcript level was unambiguously higher at 14°C than at 24°C, and following the temperature change adjusted within 30 minutes and remained similar to the control 4 hours later (Fig. 6A). Consistently, the unsaturation level in transgenics control lines was about twice as high at 14°C compared to 24°C (Fig. 6B). Overexpressing SLD8 resulted an increase of LCB Δ8-unsaturation, in particular at 24°C, where the unsaturation was comparable to EV and SLD8 at 14°C (Fig. 6B).

**Figure 6.**
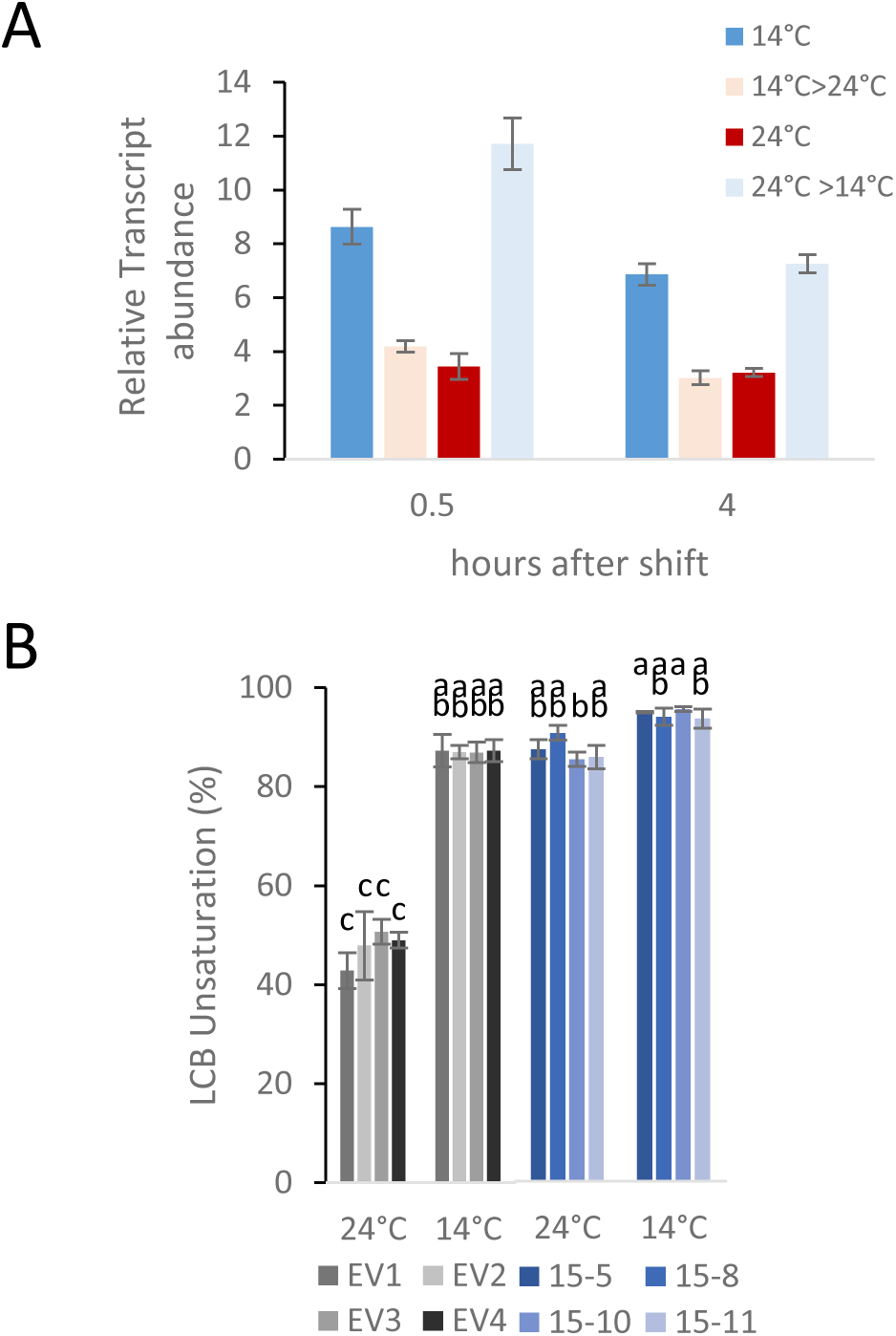
Effect of temperature on SLD8 expression and LCB unsaturation. **A**. Expression of the SLD8 according to temperature in wild-type. Exponentially growing cells were shifted from 24°C to 14°C and reversed while control cells were maintained at the initial temperatures. Cells were harvested 0.5 and 4 h after the temperature shift. Means and standard deviations of technical triplicate are shown. Experiment was repeated twice. **B.** LCB unsaturation rate in control lines (grey) and SLD8 overexpressing lines (blue). Means and standard deviations of culture triplicate are shown. Values were compared among the different transgenics according to Tukey test at p < 0.05 and groups with the same letter are not detectably different.

#### Effect of temperature on SLD8-OE growth, cell size and fluorescence

In order to gain insight into the physiological significance of SL Δ8-unsaturation according to temperature, the growth, cell size and structure as well as chlorophyll fluorescence of SLD8 OEs and negative control (EV) were monitored along a kinetics in the different temperature conditions described above (Fig. 7, Fig. S5, Fig. S6, Table S2). Cell parameters analysed by flow cytometry include two refractory parameters and chlorophyll fluorescence. Refractory parameters are FSC, (forward scatter) reflecting cell size, and SSC (side scatter) reflecting cell structure. The red fluorescence arise from the excitation of chlorophyll a at 488 nm and its emission in the red channel. Cell acclimated to 24°C and 14°C were sub-cultured and allowed to adapt for two days. Part of the culture was then switched to either lower or higher temperature 4 hours after light on in exponential growth phase (Fig. S5A-D). From our analysis, overexpression of SLD8 had no appreciable effect on growth, and EVs and SLD8-OEs displayed similar generation times under each condition. The growth was significantly enhanced at 24°C compared to 14°C and slightly slow-down upon chilling (Fig. 7A, Fig. S5E). Noteworthy, the cell parameter values reflecting size (FSC) (Fig. 7B, C), chlorophyll fluorescence (red fluorescence) (Fig. 7D, E) and cell structure/granularity (SSC) (Fig. S7 A, B) were generally higher for acclimated cells at 24°C than at 14°C. Moreover, the values in SLD8-OEs were overall lower compared to EVs, the probabilities of a significant difference being higher at 24°C than at 14°C (Fig. 7B-E, Fig. S7A-E, TableS2). In particular, the size of SLD8-OEs at 24°C was closely related to that at 14°C (Fig. 7B, C, Fig. S7E, Table S2) yielding a reduced size-ratio between 24°C and 14°C in SLD8-OES compared to EV (Fig. S7D). Interestingly, chilling allowed the size and structural differences between SLD8-OE and EV to be supressed within 24 hours after the temperature-shift. (Fig. 7C, Fig. S6B, Table S2), which is less than the time required to divide (Fig. 7A). Conversely, differences in cell-size between SLD8-OEs and EV were increased after warming though a longer period seemed to be required (Fig. 7B, Fig. S6A, Table S2). Although a similar trend occurred for chlorophyll fluorescence (Fig. 7D, E, Fig. S6D), values in cells acclimated at 14°C remained significantly lower in SLD8-OEs. Altogether, these results suggest that cell size and chlorophyll fluorescence were higher at 24°C and that overexpressing SLD8 impaired cell size adjustment at 24°C while having little effect at 14°C.

**Figure 7.**
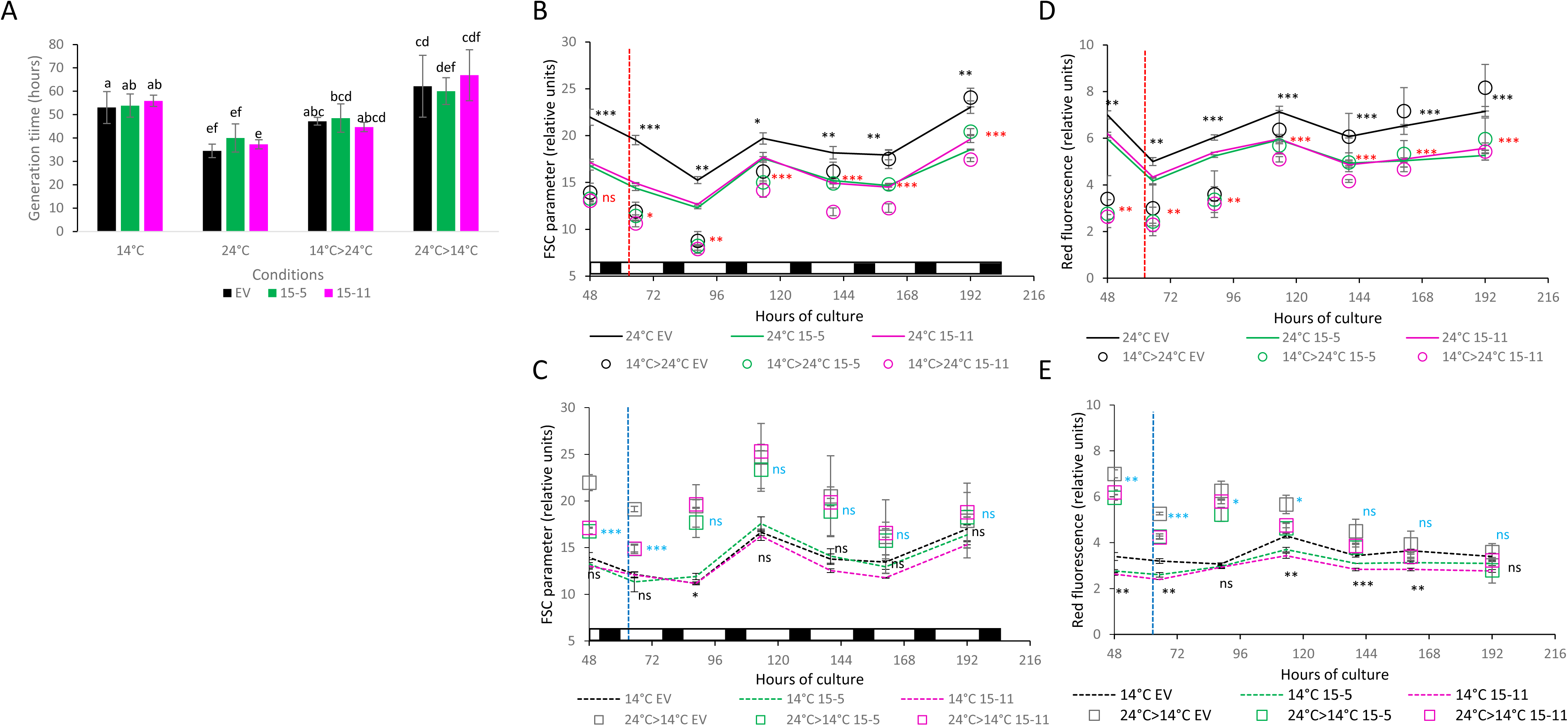
Flow cytometry analysis of growth and cell parameters of SLD8-OEs and control line under different temperature conditions. Batch cultures acclimated to 24°C or 14°C were grown under light/dark cycles (white and black boxes represented only in B and C). At 62 hours (i.e, 4 hours after light on) triplicate of the SLS8- OEs (15-5, 15-11) and negative control (empty vector, EV) were transferred from 24°C to 14°C (24°C>14°C, blue dotted line) and from 14°C to 24°C (14°C>24°C, red dotted line) and other triplicate were left at the initial temperature for control. Cell counts, refractory parameters (FSC) and red fluorescence parameters were monitored along the growing kinetics. **A**. Generation times during exponential growth after temperature shift. Values were compared among the different transgenics according to Tukey test at p < 0.05. Paiwise t-test within each conditions yield p > 0.1 (ns) (Table S2). Evolution of cell size (FSC) at 24°C (**B**) and 14°C (**C**). Evolution of chlorophyll-a fluorescence at 24°C (**D**) and 14°C (**E**). means an SD are shown Statistical significance by t-test. (*, p < 0.1; **, p < 0.05; ***, p < 0.001; ns, not significant). For clarity, the statistical values represented are for 15-5 compared to EVs at each time point for acclimated cells (black) and cell shifted to lower (blue) or higher temperature (red). See Fig . S5 for growth curves, Fig S6 for SSC parameter and supplemental graphs for FSC and fluorescence parameters. Complete statistical values are available from Table S2.

### Characterisation of *O. tauri* sphingolipids

To further gain insight into the SLs substrate of SLD8 and possibly involved in cell-size adjustment defect, the detailed characterization of *O. tauri* SL was first carried out in wild type (Fig. 8, Fig. S7). In land plants and some algae, GIPC and GlcCer are known as the major SLs (Mamode Cassim et al., 2020). GlycoSLs provide the specific product ions (i.e., ceramide and LCB moieties) in MS/MS analysis under a positive ionization mode. Based on the above mentioned ceramide composition of *O. tauri* (Fig. 4), the total lipid extract from *O. tauri* was applied to the precursor ion scanning using the fragment of d18:1 LCB (*m*/*z* 264.3, [LCB−2H_2_O+H]^+^) and d18:1-16:0 ceramide (*m*/*z* 520.5, [Cer−H_2_O+H]^+^) (Fig. 8, Table S1). The MS^2^ analysis provided the *m*/*z* of putative precursor species 538.5, 714.6 and 890.6 for LCB (Fig 8A) and 714.5, 890.6, 1052.6, 1214.7, and 1376.7 for Cer (Fig 8B). The *m*/*z* 538.5 corresponds to d18:1-c16:0 Cer, and thus the larger fragments were estimated as glycoSLs. The differences between the *m*/*z* 890.6, 1052.6, 1214.7 and 1376.7 are Δ162, which can be unambiguously attributed to hexose (Hex) attachments (Fig. 8C). There are therefore up to three Hex residues at the terminal end of SLs. In addition, the differences Δ176 (*m*/*z* 890.6 from 714.6 and *m*/*z* 714.6 from 538.5) and Δ194 (*m*/z 714.6 from 520.5) can be attributed to hexuronic acid (HexA, without or with H_2_O, respectively). Thereof, up to two HexA residues are attached to the base of the glycosyl chain (Fig. 8C). Vice Versa, the product ion scanning of the predicted precursor ions supported the glycan structure (Fig S7). However, highly glycosylated molecules are often difficult to be unambiguously determined by ESI-MS, as multiple deglycosylated fragments are detected via in-source fragmentation. We therefore assessed the above results by further performing targeted MS^2^ analysis coupled with a reverse-phase HPLC separation. This enabled us to detect four separate peaks in the LC chromatogram, which were assigned to Cer, HexACer, Hex_2_HexA_2_Cer, and Hex_3_HexA_2_Cer (Fig. 8D). The chromatograms obtained with MRM settings for HexA_2_Cer and HexHexA_2_Cer were overlapped with that for Hex_2_HexA_2_Cer, and the peak at 9.7 min of retention time was also detected in the windows for Cer and HexACer but not for Hex_3_HexA_2_Cer where a distinct peak was detected at 9.2 min, indicating that the overlapped peaks are Hex_2_HexA_2_Cer and its in-source fragmentation products. Note that neither (G)IPC nor GlcCers were detected by the scanning and targeted analysis that has been successively developed for the SLs in land plants (Ishikawa et al., 2016). *O. tauri* therefore encompasses peculiar classes of complex and acidic glycoSLs ranging to a simple monoglycosylSLs to pentaglycosylSLs. Most importantly, the basis of the glycosyl chain consist of two hexuronic acids. GlyCers composed of hexuronic acids have so far only been identified from the proteobacteria genus Sphingomonas (Kawahara et al., 2000).

**Figure 8.**
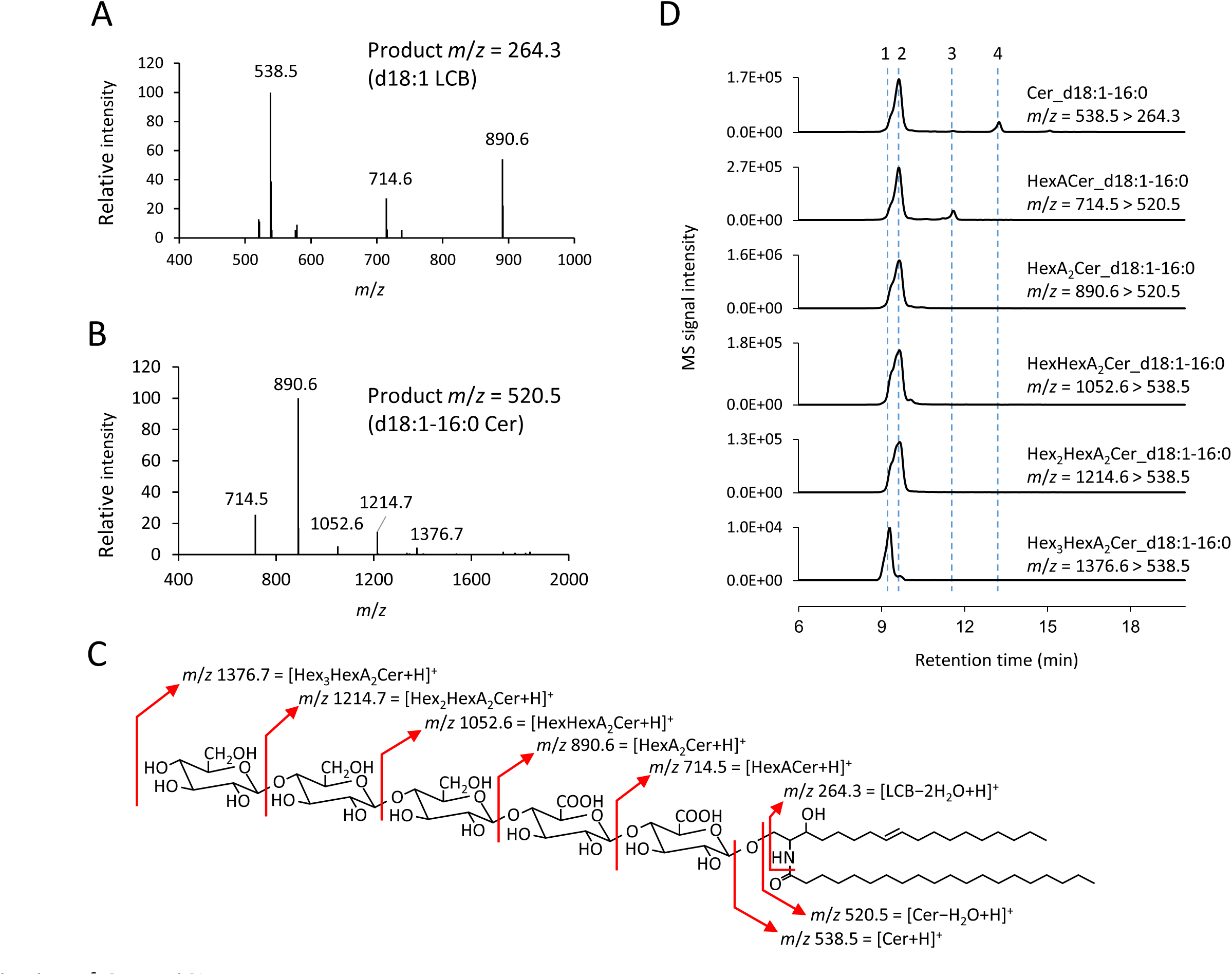
Characterisation of *O*. *tauri* SLs. **A**. Precursor ion scanning using the product ion corresponding to d18:1 LCB. **B**. Precursor ion scanning using the product ion corresponding to d18:1-16:0 Cer. **C**. Assignment of the detected *m*/*z* to glycosphingolipids containing HexA and Hex residues. **D**. Targeted LC-MS/MS chromatogram of *O*. *tauri* SLs. Four separate peaks were determined as below; 1, Hex_3_HexA_2_Cer; 2, Hex_2_HexA_2_Cer; 3, HexACer; 4, free Cer. The MRMs used were combination of molecular ions ([M+H]^+^) and the major product ions for each species, i.e., *m*/*z* 264.3 ([d18:1-2H_2_O+H]^+^) for Cer, 520.5 ([Cer-H_2_O+H]^+^) for HexACer, and 538.5 ([Cer+H]^+^) for longer glycoSLs as determined by the product ion scanning (Fig S7).

### Effect of temperature and SLD8 overexpression on sphingolipids

The next step was to obtain the relative class composition of SL and to determine the possible influence of temperature and of SLD8 overexpression on these classes. To this end detailed SL analysis of SLD8 overexpressors and EVs acclimated at 14°C and 24°C was carried out (Fig. 9, Fig. S8). Individual changes were analysed in each of the four EVs and SLD8-OE (Fig. 9A, B) and a general trend was gained by averaging all values for each line type (Fig. 9C- F). Overall temperature and *SLD8* overexpression had a major impact on the SL unsaturation while the impact on the amount was more subtle. The most abundant (as signal intensity) SL classes were the mono-GlyCer, HexACer and the tetra-GlyCer, Hex_2_HexA_2_Cer, while Cer and Hex_3_HexA_2_Cer were detected at lower intensities (Fig. 9A-C). Comparing 14°C to 24°C, a general trend consisted of a reduction of Hex_2_HexACer and of Cer, and of an increase of HexAcer and Hex_3_HeXA_2_Cer (Fig. 9C). These changes occurred to a relative similar extent in OEs and EVs, although the amount of Hex_2_HexA_2_Cer in SLD8-OEs remained high at 14°C (Fig.9C, 9D). All SLs displayed Δ8-unsaturation, the most complex SLs Hex_3_HeXA_2_Cer being the most unsaturated, followed by Hex_2_HexA_2_Cer and then Cer, whereas HexACer were the least unsaturated SLs (Fig.9E). The degree of unsaturation was higher in all SL classes at 14°C in EVs but also in SLD8-OEs, which already unambiguously displayed a higher unsaturation compared to control lines at 24°C (Fig. 9A, B, E). In particular, the unsaturation degree in SLD8-OEs at 24°C appeared closely related to that of EVs at 14°C (Fig. 9E). It should be noted that the unsaturation in SLD8-OE at 14°C is even higher. To summarize it appears that 1) HexACer and Hex2HexA2Cer are the most abundant SLs, HexACer production being favoured at low temperature, 2) low temperature as well as SLD8 overexpression boost the unsaturation in all SL classes.

**Figure 9.**
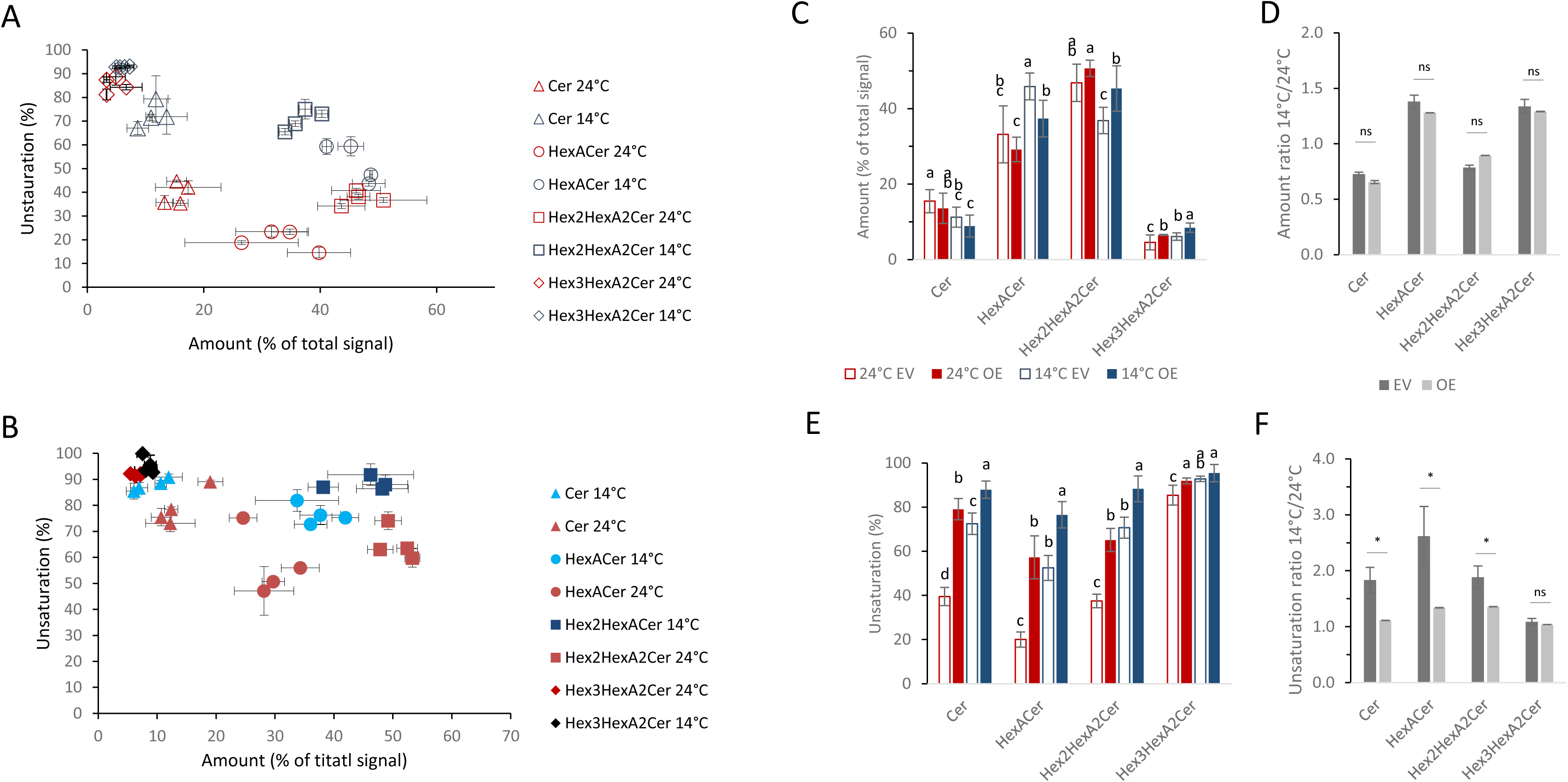
Sphingolipid contents and unsaturation rates in control and SLD8 overexpressing lines according to temperature. Four individual lines where cultivated in triplicate for each the negative control (EV1,EV2,EV3,EV4, open colored symbols) and SLD8-OEs (15-5, 15-8, 15-10, 15-11 plain colored symbols). Cells acclimated at 24°C and 14°C were collected at the end of exponential growth for analysis. Correlation between sphingolipid unsaturation and relative content in individual EVs (**A**) and SLD8-OEs (**B**). Means and standard deviations of triplicate culture are shown. **C**-**F**. All values for each line type EV (n=12) and SLD8-OE (n=12) were averaged. **C**. Relative amount of sphingolipids in EVs and SLD8-OEs at 14°C and 24°C. **D**. Ratio of relative amount between 14°C and 24°C. **E**. Unsaturation rate in sphingolipid classes in EVs and SLD8-OEs at 14°C and 24°C. The following formula was used d18:1/(d18:0+d18:1) ×100 (%). F. Unsaturation ratio between 14°C and 24°C. Stastistical significances within each SL class are shown as different letters in C and E (Tukey test, p < 0.05) and as an asterisk in D and E (Student’s t-test, p < 0.05, ns = not significant). Errors bars for ratio z=x/y in D and F were calculated according to the error propagation rule 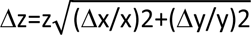.

### Sphingolipid genes

Based on our results and further retrieval of *O. tauri* SLs related genes homologues, we propose a SL pathway in *O. tauri* (Fig. 10, Fig. S9). Concerning LCB/ceramide modification enzymes, no SLD4 orthologue could be retrieved from the *O. tauri* genome, though putative homologues were found in *Micromonas* species. No homologue of fatty-acid hydroxylase (AtFAH1) occurred in any Mamiellales species but *B. prasinos*. These results are consistent with our analysis which shows that apart from LCB Δ8-unsaturation, no other changes are detected. However, although we did not detect a t18:0, a putative homologue of LCB C4- hydroxylase was found in *Ostreococcus* species and *Micromonas commoda*. Surprisingly, a putative homologue of the LCB C9-methyltransferase, usually found in fungi, also occurred in all Mamiellales species. Putative LCB/ceramide-P phosphatase homologues were also retrieved.

**Fig. 10.**
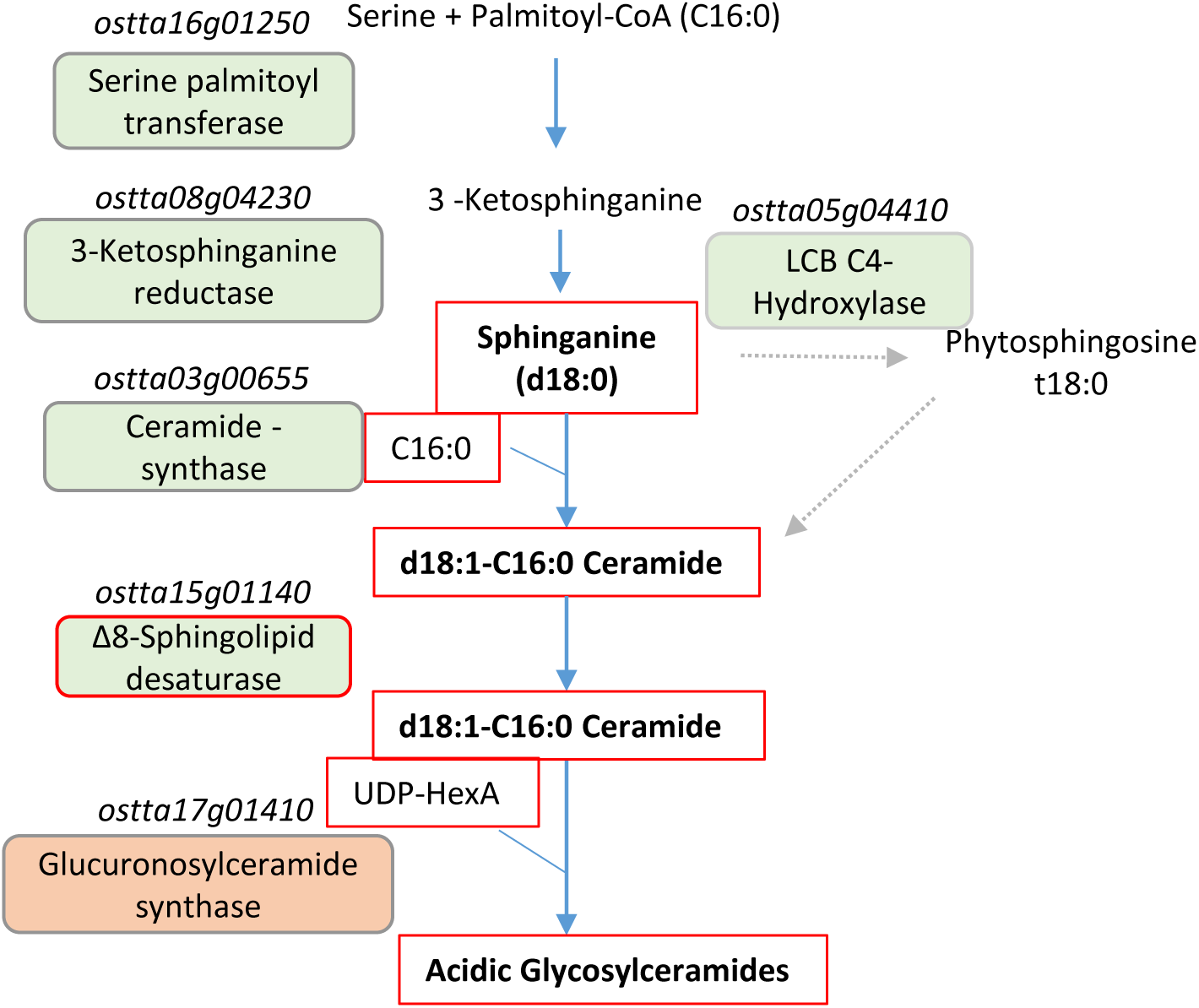
Sphingolipid pathway in *O. tauri*. Red frames highlight sphingolipid and enzyme characterised in the present work. Enzymes involved in the metabolic step are indicated and corresponding genomic accession are indicated (italics). Homologues with E-values ≤ 9.00E-07 and query cover ≥ 41% are indicated in green boxes when obtained by blasting *A. thaliana* protein accessions and in pink box when obtained by blasting protein accessions from other organisms. Grey arrows represent unvalidated step. Details about blast results are provided in Fig. S9.

Core enzymes for SL synthesis readily identified encompassed the serine palmitoyl transferase (SPT) and ceramide synthase, which is more closely related to CS-like from the SAR lineage. As regards the glycosylceramide synthase (GSC), putative *A. thaliana* homologue could be retrieved from *Bathycoccus* and *Micromonas* but not from *Ostreococus* species. However, putative homologues of the recently characterised glucuronosylceramide synthase from the alpha proteobacterium *Zymomonas mobilis* occured in *O. tauri* as well as in other Mamiellales species (Okino et al., 2020). In agreement with our analysis, the search for enzymes involved in the synthesis of GIPC (IPC-synthase, inositolphosphoryl ceramide synthase) proved unsuccessful.

## DISCUSSION

More than twenty-five years ago, Sperling and collaborators cloned from plant, the first SLD8 desaturase opening the way for investigating the evolution and physiological functions of SL Δ8-unsaturation (Sperling et al., 1998). Compared to plants, very little data are available about the structure and function of SLs in microalgae. Once again, it is important to emphasise that microalgae are distributed in four supergroups of eukaryotes and represent a much greater biological diversity than plants (Archaepalstida supergroup). *O. tauri*, occupies a basal position in the green lineage which makes this unicellular organism key to elucidating early stages in the history of lipid evolution. Previous detailed glycerolipidome characterization of *O. tauri* unveiled features at the cross between SAR and Archaeplastida. In the present study we not only provide the first evidence of the involvement of SL Δ8-unsaturation in temperature acclimation of microalgae, but also report novel eukaryotic SLs features that might have been acquired early in the evolution.

### Main *O. tauri* SLs display unique feature reminiscent of bacteria glycosylceramides

From the few reports on microalgal SLs, the first structural insight into microalgal GlyCers have been acquired from the green microalga *Tetraselmis* sp., which is related though paraphyletic, to *Ostreococcus* (group previously referred to as Prasinophyceae) (Arakaki et al., 2013). *Tetraselmis* ceramides glycosyl moiety corresponds to a single glucose residue. From structural analyses of SLs of 17 strains of the SAR supergroup it was shown that GlyCer consisted of one to three monosaccharide units linked to highly diverse ceramides backbones (Li et al., 2017). Our work unveils that *O. tauri* displays three unique classes of GlyCer consisting of hexuronic acids at the basis of the glycosyl head, a feature that makes them unique as hexuronic acid has only been reported from bacterial SLs (Kawahara et al., 2000; Vítová et al., 2022). In particular, glucuronic acid or galacturonic acid residues α-linked to sphingosine derivatives were reported from the proteobacteria *Sphingomonas*. The GlyCer are structurally related to lipopolysacharrides which are absent in sphingomonads. In *O. tauri,* two hexuronic acids make the basis of the glycosyl chain consisting of up to three additional hexosyl residues. Hex_2_HexA_2_Cer and HexACer are the major SLs class, the former prevailing at 24°C and the latter at 14°C. Consistently, putative orthologues of glucuronylceramide synthase from the alpha proteobacterium *Z. mobilis* are present in Mamiellales species (Okino et al., 2020). While beyond the scope of the present study, it would be interesting to assess the specificity of these enzymes in the future.

Like in bacteria or fungi, the ceramide backbones of *O. tauri* SLs consist of d18:0/d18:1 and C16:0 with C14:0 occurring in trace amount. Although phytosphingosine has been sporadically reported from few green microalgae, it is generally absent in most microalgae and *O. tauri* is no exception (Shiels et al., 2021). However, it is more surprising that *O. tauri* ceramides do not contain long-chain acyl LCBs or polyunsaturated LCBs. Indeed, other marine species such as *Tetraselmis* sp. and diatoms that share with *O. tauri* the ability to synthesise VLC-PUFAs display such characteristics (Arakaki et al., 2013; Li et al., 2017).

### *O. tauri* SLD8 sequence features and substrate specificity shed light on SLD8 evolution

The evolution of eukaryotic desaturases, in particular of CytB5-fused desaturases has been thoroughly investigated in the past, and hypotheses about the evolution of desaturase substrate specificity have been proposed (López Alonso et al., 2003; Sperling et al., 2003; Gostincar et al., 2010). It is worth recalling here that SLD8s only occur in plants and fungi, and further that plant Δ6-Des only occur in few 18:3n-6 producing species, uses lipids as substrates whereas Δ6-Des are widespread in animals but uses acyl-CoAs as substrates. *O. tauri* related species are therefore unique in the plant lineage as, in addition to SLD8, they display an animal-like acyl-CoA Δ6-Des (ER) and plastidic Δ6-Des that use lipids as substrates (Domergue et al., 2005; Degraeve-Guilbault et al., 2020). Note, however, that previous phylogenetic analyses revealed that Mamiellales acyl-CoA-Δ6-Des, branch separately from their animals counterparts (Gostincar et al., 2010).

From our phylogenetic analysis, SLD8, pΔ6-Des and acyl-CoA-Δ6-Des clades of Mamiellales species are robustly related, with SLD8 and pΔ6-Des clustering together while acyl-CoA Δ6-Des clade sets more apart. On the other hand, OtSLD8 robustly clusters with plant SLD8. Sperling proposed that “*a common and ancient fusion desaturase with Δ5- or Δ6- regioselectivity* [is at the origin of CytB5-fused desaturase]. *Comparatively late gene duplications gave rise to “local”, i.e. parallel developments of some of the basic regioselectivities* [including] *Δ6 from sphingolipid-Δ8* (plants)” (Sperling et al., 2003). Accordingly to Gostincar and collaborators, the duplications and specializations of Δ6/Δ8 pairs have happened more than once (Gostincar et al., 2010). Our analysis suggests a common ancestor to acyl-CoA-Δ6-Des, plastidic Δ6-Des and SLD8. Either the ancestor desaturase had a Δ6-regioselectivity and gave rise to SLD8 after later gene duplication or the ancestor had a mixed specificity and regioselectivity specifications occurred after gene duplications in parallel to give rise to pΔ6-Des, SLD8 and acyl-CoA Δ6-Des.

OtSLD8, displayed a clear preference for dihydroxylated substrates, whether they were saturated or Δ4-unsaturated, and produced exclusively *E*-isomers. These features are strongly reminiscent of the distantly diatom SLD8 homologue TpdesB (Tonon et al., 2005). It is worth recalling that SL-Δ4 desaturation is more widespread in eukaryotes than Δ8 desaturation and is regarded as an ancestral feature within the green lineage (Haslam and Feussner, 2022). Moreover, experiments in yeast and moss suggested that Δ4 unsaturation precedes, and is required for, Δ8 desaturation (Ternes et al., 2011; Gömann et al., 2021). Our results indicate that this is not the case for OtSLD8. Furthermore, OtSLD8 activity on trihydroxylated is very poor, which contrasts with AtSLD1 but recalls of TpdesB. Although functional characterisation of additional microalgal homologues are required to gain insight into the evolution of SLD8 specificities, it is tempting to speculate that the ancestral specificity of SLD8 is not dependent on LCB C4 modifications. Further reasoning that microalgae from the SAR and basal to Archaplastidia have conserved the ancient substrate specificities of plant SLD8, the specificities for C-4 modified LCB would have been acquired later in the evolution. An alternative is the loss of LCB Δ4 unsaturation requirement for SLD8 activity in *T. pseudonana*, *O. tauri* and *A. thaliana*. Nevertheless, it seems reasonable to speculate that the *E*-regiospecificity occurring in fungi, diatoms and *O. tauri*, is an ancestral feature, inherited very early in the evolution; the *Z*-isomer synthesis would rather be a shared derived character of land plants (synapomorphy).

Finally, the occurrence of a cTP in SLD8 sequences from Mamiellales and the plastidic localization of the long-OtSLD8 in *N. benthamiana* is puzzling. Indeed, it is commonly assumed that SLD8 are ER enzymes. Nevertheless, links between SLs and chloroplasts exists and recent data showed that tomato plants silenced for *SLD8* and exposed to chilling displayed severely damaged chloroplasts (Zhou et al., 2016). In the case of *O. tauri*, the possibility of the dual localization (ER/plastid) of some desaturases including SLD8 remains in doubt and has to wait further technical developments allowing subcellular localisation in the tiny native host. Nevertheless, the occurrence of a second methionine upstream of the annotated ORF is common to several *O. tauri* desaturases including the plastidic ω3-Des, we recently characterised (Degraeve-Guilbault et al., 2021). As a ω3-Des activity is also required outside the chloroplast and no other sequence is available in Mamiellales, we had proposed that alternative translation could allow the synthesis of two different isoforms of the ω3- Des. Alternative translation could possibly be also at work for producing two differentially located SLD8. Finally, it should be recalled that *O. tauri* is an ultra-compact organism in which the ER is extremely reduced while the nuclear envelope displays large extensions in contact with the chloroplast (Henderson et al., 2007). Thus, plastidic enzymes could possibly have access to extra-plastidic lipids through contact sites with other organelles.

### Physiological function of OtSLD8

Reports about physiological function of SLs in algae are scarce: SLs are believed to be crucial component of the interaction between *Emiliania huxleyi* and its cognate virus (Zeng et al., 2019). SLs have also been shown to be involved in ciliary motility in the microalgae *Chlamydomonas reinhardtii* (Kong et al., 2015). However, whether desaturation of SLs is involved remains elusive. Insights into the physiological role of SLs unsaturation and of SLs Δ8-unsaturation in particular, has been mostly gained from plants and mostly from *A. thaliana*. Chen and co-workers demonstrated that the *Arabidopsis* double *sld1 sld2* mutant which completely lacks SL Δ8-unsaturation, failed to acclimate to low temperature (Chen et al., 2012). Accordingly, *sld1* and *sld2* transcript expression increased at low temperature (Nagano et al., 2014). Moreover, lipidomic analyses of natural *Arabidopsis* accessions more or less resistant to low temperature showed that the relative content of the unsaturated GlcCer d18:1,C24:0 was positively correlated with increased freezing tolerance (Degenkolbe et al., 2012). Finally, Zhou and co-worker showed that in tomato *SlSLD* knockdown plants, the ultrastructure of envelope was severely affected after exposure to chilling stress (Zhou et al., 2016).

Our work shows that in *O. tauri* temperature regulates Ot*SLD8* expression and SL Δ8- content and unsaturation as well as cell size and natural fluorescence. SLD8 transcript and SL Δ8-unsaturation are increased at 14°C, temperature at which cell size, cell granularity and natural fluorescence are decreased compared to 24°C. Overexpression of SLD8 suppresses the decrease in transcription at 24°C and results in an increase in SL-Δ8 unsaturation to a level similar to that of control lines at 14°C. Cell growth of SLD8 OEs remains unaffected; SLD8 OEs however fail to adjust their size and natural fluorescence at 24°C. SLD8-OEs display clear defect in the regulation of cell size at 24°C and not at 14°C, which is consistent with the marked increase of SL-unsaturation at 24°C. Since in SLD8-OEs the unsaturation degree is higher in all SLs and only minor changes occur in the content of major classes, it is difficult to speculate about the SLs class that would be possibly involved. Chlorophyll content a well as chlorophyll fluorescence decrease have been reported in chilled plants, and related to photosynthesis adaptation (Riva-Roveda et al., 2016; Daems et al., 2022). Since the chloroplast occupies most of the cell volume in *O. tauri* the variation in size might to some extent be related to the chloroplast adaptation to temperature (Henderson et al., 2007). Independently of the physiological significance of these changes, our results unambiguously indicate that the downregulation of SLD8 at high temperature is required for *O. tauri* cell-size and/or chlorophyll fluorescence adaptation. The biological significance of these changes and the involvement of sphingolipids will deserve further investigations.

## METHODS

### Chemicals

Chemicals were purchased from Merck, Sigma Chemical (St. Louis, MO, USA) and Wako (Tokyo, Japan) when not otherwise stated. Lipids used as internal standards were purchased from Avanti Polar Lipids (Alabaster, AL, USA). Internal standards were prepared according to Markham et al. (2007).

### Sequences and phylogeny analyses

Sequences were obtained from NCBI. SLD8 from Mamiellophyceae species were manually checked for completion of Nt sequences as reported previously for other desaturases (Degraeve-Guilbault et al., 2020; Degraeve-Guilbault et al., 2021). Briefly, open reading frame (ORF) encompassing a start methionine upstream of the annotated start codon were taken into account. Target peptides were predicted from PredAlgo (Tardif et al., 2012). Desaturase sequences alignment was performed using Snapgene trial version (Clustal omega).

Phylogenetic analyses used OtSLD8 as query to retrieve homologs from as many taxa as possible. The first dataset produced contained hundreds of sequences with the most populated phylum in plants. A subset of plant sequences was chosen and encompassed sequences of proteins that has been functionally characterised (Li et al., 2016). The subset was then back-BLASTed in all the major repositories (NCBI, Ensembl, JGI) and a new dataset containing SAR and fungal protein was produced. The dataset was visualized in BioEdit v7.0.5.3 computer program then aligned and curated using the MAFFT v7.407_1 (Katoh and Standley, 2013)and BMGE v1.12_1 (Criscuolo and Gribaldo, 2010) tools implemented in NGphylogeny.fr (Lemoine et al., 2019). The final alignment included 46 sequences and 326 residues including gaps. Two independent Metropolis-Coupled Markov chain Monte Carlo (MCMCMC) algorithm was used to infer Bayesian phylogenetic relationships using MrBayes v3.2.7_0 software (Ronquist et al., 2012) implemented in NGphylogeny.fr. Each analysis involved one cold and three warm chains. The number of generations was set at 2.000.000 and with a sampling frequency every 500 generations and a burn-in fraction set at 0.25, i.e. the first 500.000 tree topologies were discarded to stabilize the analyses. The parameters of the likelihood model was set to ‘mixed’ hence the algorithm was allowed to jump from one model implemented in the software to the other according to dataset. A gamma distribution probability distribution was set. Once the analysis completed, the outcome was saved in the Nexus format and then visualized in FigTree v1.4.4 to set colors.

The histidine boxes previously identified were aligned following the clades produced in the Bayesian inference and were used to feed WebLogo3 (Crooks et al., 2004) online tool to produce the sequence logos in Figure 1.

### Biological material and cultures

*O. tauri* cells (clonal isolate from OtH95) were grown under 18h light- 6 h dark cycles in artificial sea-water (ASW) NaCl 4.1 ×10^-1^ M, KCL 8×10^-3^ M, MgCl_2_.6H_2_O 3.2×10^-2^ M, CaCl_2_ 2.7×10^-3^ M, Tris –HCl 5mM pH 7.6 supplemented with F/2 components without silicium (ncma.bigelow.org/algal-recipes). Cultures were grown in incubator-shaker (New Brunswick Innova 42R) with constant agitation (80 RPM) under white light (75 mmol photons m^-2^ s^-1^, 6 X T8 fluorescent bulbs 15 Watt each (Sylvania Gro-Lux). For screening of FA of *O. tauri* transgenics, cells were grown in T25 aerated culture flasks (Sartstedt, Nümbrecht, Germany) at 20°C. Phosphate (NaH_2_PO_4_) was reduced from 35 µM to 5 µM to allow the expression of transgenes under the high affinity phosphate transporter promoter. Bacteria associated to *O. tauri* were reduced to less than 5% using centrifugation (1500 g, 5 min, RT) and antibiotic treatment (vancomycin 1 mg/mL, 4 days followed by G418 0.5 mg/ml, 4 days). Survey of bacterial population was achieved by flow cytometry as previously reported (Degraeve-Guilbault et al., 2017).

### Flow cytometry

Flow cytometry (Partec CyFlow Space FACS Flow Cytometer) analyses were achieved on fixed cells (10 min, 0.5 % glutaraldehyde Grade II). The light scattered parameters forward scatter (FSC) and side scatter (SSC) were used for the discrimination of cells by size and cell internal complexity (i.e granularity), respectively. *O. tauri* cell were discriminated on the basis of the red fluorescence and counted.

### Temperature shift

Cell were acclimated to 24°C and 14°C for at least 3 weeks (3 subculturing) during which the cell density strains was regularly adjusted in order to remain similar between conditions and transgenics and low phosphate medium was used in the last round of subculturing to allow the full expression of transgenes. By the time 0 of the experiment, each batch was subcultured in 6 independent flasks and allowed to recover for 62 hours before the temperature shift. Sampling of 1 mL of culture was achieved along the growth kinetics and fixed cells were analysed within 6 h.

### Cloning strategy

PCR amplifications of DES ORF were achieved using Q5® Polymerase by two-step PCR on cDNA matrix (5’-GGGCCCATGGCGTCGTCGGTGGGCG ATGCGCGCCGCGACGTC-3’.and 5’-CTAGTCGCCCCGCTCCCAGAC-3’), which encompassed the restriction sites ApaI for cloning in pOtox-Luc (Moulager et al., 2010). The amplified fragment was subcloned in pGEM®-T Easy (Promega, Madison, WI, US) and sequenced (Genwiz, Leipzig, Germany). Cloning using Gateway® system was performed according to manufacturer instruction (pDONR 221, pK7W2G2D or pK7YWG2 for N. benthamiana). Sequences for either the long or the short version of SLD8 were amplified from the SLD8 sequence cloned in pGEMT with primer encompassing the extension AttB1 and AttB2 extension. Codon optimized sequences (GenScript Biotech, Netherlands) were used for overexpression in S. cerevisiae.

### RNA and cDNA preparation and quantitative RT-PCR analysis

RNeasy-Plus Mini kit (Qiagen, Hilden, Germany) was used for RNA purification; DNase I was used to remove contaminating DNA (DNA-free kit, Invitrogen, Carlsbad, USA) and cDNA obtained using the reverse transcription iScript^TM^ supermix kit (Bio-Rad, Hercules, CA, USA). Primers used for qPCR were (5’-CCCTTCGCGGAAAAGAATGG-3’.and 5’- GCTTGAGCGTTCGAAACACC-3’). Real-time RT quantitative PCR reactions were performed in a CFX96^TM^ Real-Time System (Bio-Rad) using the GoTaq^®^ qPCR Master mix (Promega, Madison, WI, USA). Bio-Rad CFX Manager software was used for data acquisition and analysis (version 3.1, Bio-Rad). Ct method was used to normalize transcript abundance with the references *ACTprot2* (Actin protein-related 2). PCR efficiency ranged from 95 to 105%. Technical triplicate were used, and at least two independent experiments were achieved.

### Genetic transformation

*O. tauri* electroporation was adapted from (Corellou et al., 2009). Transgenics were obtained by electroporation and pre-screened accordingly to their luminescent level (Degraeve-Guilbault et al., 2020). *S. cerevisiae* was transformed by the standard LiAc/SS carrier DNA/PEG method (Gietz and Schiestl, 2007) using pEG423 vector. Control lines are transgenics of empty vectors. *N. benthamiana* leaves from five-week old plants were infiltrated with *Agrobacterium tumefaciens* previously transformed by electroporation; the p19 protein to minimize plant post-transcriptional gene silencing (PTGS) was used in all experiments (Shah et al., 2013). Briefly, *A. tumefaciens* transformants were selected with antibiotics (gentamycin 25 µg/mL with spectinomycin 100 µg/mL or kanamycin 50 µg/mL). *Agrobacterium* transformants were grown overnight, diluted to an OD_600_ of 0.1 and grown to an OD_600_ of 0.6-0.8. Cells were re-suspended in 5 mL sterilized water for a final OD_600_ of 0.4 and 0.2 for overexpression and subcellular localization experiments, respectively and 1 mL was agroinfiltrated. Plants were analysed 2 and 5 days after *Agrobacterium* infiltration for subcellular localization experiments and for overexpression, respectively.

### Lipid analysis

For all organisms, FA analyses and for *O. tauri* further lipid analysis were achieved as previously reported (Degraeve-Guilbault et al., 2017).

To obtain total LCB composition, lyophilized *O. tauri* cells or fresh yeast cells were directly hydrolyzed in 10% barium hydroxide/dioxane (1:1) for 24 h at 110°C. After cooling, the solvent was vigorously mixed with 0.5 vol of diether ether and 4% sodium sulfate and the upper phase was collected and evaporated. LCB was derivatized with NBD-F and analyzed by LC-MS/MS according to the previous report (Ishikawa et al., 2014) with the MRMs shown in Table S1.

Total sphingolipid extract was prepared and analyzed as previously described with some modifications (Ishikawa et al., 2016; Ukawa et al., 2022). In brief, lyophilized cells were homogenized by vigorous shaking with glass beads in 1-butanol/methanol (2:1). The homogenate was mixed with 0.6 vol of 1 N KOH and kept for 30 min at 50°C. The mixture was acidified with HCl and phase-separated by adding 1-butanol. The upper layer was collected and dried in vacuo. The residue was dissolved in THF/methanol/water containing 0.1% formic acid and injected to LC-MS/MS (LCMS-8030, Shimadzu GLC). For MS/MS analysis of glycoSL structures, the sample was directly injected into MS under a flow of methanol/isopropanol/acetonitrile (1:1:1) containing 5 mM ammonium formate. Precursor ion scanning analysis was performed with product ion of m/z 264.3 (LCB) or 520.5 (Cer) and a range of precursor ion scan between *m*/*z* 250 to 1800. Targeted MS/MS analysis was performed using the MRMs shown in Table S1. The solvent system and MS parameters were according to Ishikawa et al. (2016).

### Confocal microscopy

Live cell imaging was performed using a Leica SP5 confocal laser scanning microscopy system (Leica, Wetzlar, Germany) equipped with Argon, DPSS, He-Ne lasers, hybrid detectors, and 63x oil-immersion objective. *N. benthamiana* leaf samples were transferred between a glass slide and coverslip in a drop of water. Fluorescence was collected using excitation/emission wavelengths of 488/490-540 nm for chlorophyll, 488/575- 610 nm for YFP. Co-localisation images were taken using sequential scanning between frames. Experiments were performed using strictly identical confocal acquisition parameters (*e.g.* laser power, gain, zoom factor, resolution, and emission wavelengths reception), with detector settings optimized for low background and no pixel saturation.

### Stastistics

Statistics were performed with Excel. Student’s test was applied for comparing two data-sets.Tukey test was applied for comparing more than two data sets at once (multiple pair-wise comparison, p-value < 0.05). Compact letter display was used to clarify the output. Each variable that shares a mean that is not statistically different from another one will share the same letter. (Schlattmann and Dirnagl, 2010). Groups with the same letter are not detectably different (are in the same set) and groups that are detectably different get different letters (different sets). Groups that have more than one letter reflect “overlap” between the sets of groups.

The wild-type strain (W303-1A), sur2Δ (BY4741), and sur2Δ expressing the Δ4 sphingolipid desaturase of Komagataella pastoris were used to determine the desaturation activity on t18:0, d18:0, and d18:14E, respectively. The percent of unsaturation was calculated by the formula (product)/(substrate + product)*100. Means and standard deviations are shown (n=3).*NE: not examined due to the same Pro-GAL1 plasmid used for expression of OtSLD8/AtSLD1 and Pichia Δ4 desaturase.

## ACKNOWLEDGMENTS AND FUNDINGS

Routine lipid analyses were performed at the Metabolome Facility of Bordeaux-MetaboHUB (ANR-11-INBS-0010). Imaging was performed at the Bordeaux Imaging Center, member of the national infrastructure France BioImaging. The yeast expression vector pGK423 was provided by the National Bioresource Project (NBRP, Japan).

## AUTHOR CONTRIBUTIONS

AA and FC performed the phylogenetic analyses. FC performed most of the experimental work and analyses) related to *O. tauri* (cultures, cytometry analyses, cloning, transformation, FAMES analysis, RT-qPCR) and wrote the MS. FD performed the work and analyses on *N. benthamiana* (cloning, agro-transformation, and FAMES analysis). TI performed the experimental work and analyses related to desaturase expression in yeast and *O. tauri* sphingolipid characterisation and wrote the corresponding sections. All authors contributed to the article and approved the submitted version.

## REFERENCES

Ali U, Li H, Wang X, Guo L (2018) Emerging Roles of Sphingolipid Signaling in Plant Response to Biotic and Abiotic Stresses. Mol Plant 11: 1328–1343

Arakaki A, Iwama D, Liang Y, Murakami N, Ishikura M, Tanaka T, Matsunaga T (2013) Glycosylceramides from marine green microalga Tetraselmis sp. Phytochemistry 85: 107–114

Breslow DK, Weissman JS (2010) Membranes in balance: mechanisms of sphingolipid homeostasis. Molecular cell 40: 267–279

Cacas J-L, Buré C, Furt F, Maalouf J-P, Badoc A, Cluzet S, Schmitter J-M, Antajan E, Mongrand S (2013) Biochemical survey of the polar head of plant glycosylinositolphosphoceramides unravels broad diversity. Phytochemistry 96: 191–200

Cacas JL, Furt F, Le Guedard M, Schmitter JM, Bure C, Gerbeau-Pissot P, Moreau P, Bessoule JJ, Simon-Plas F, Mongrand S (2012) Lipids of plant membrane rafts. Prog Lipid Res 51: 272–299

Chen M, Markham JE, Cahoon EB (2012) Sphingolipid Δ8 unsaturation is important for glucosylceramide biosynthesis and low-temperature performance in Arabidopsis. Plant J 69: 769–781

Courties C, Vaquer A, RTrousselier M, Lautier J, Chrétiennot-Dinet M-J, Neveux J, Machado C (1994) Smallest eukaryotic organism. Nature 370: 255

Criscuolo A, Gribaldo S (2010) BMGE (Block Mapping and Gathering with Entropy): a new software for selection of phylogenetic informative regions from multiple sequence alignments. BMC Evolutionary Biology 10: 210

Crooks GE, Hon G, Chandonia JM, Brenner SE (2004) WebLogo: a sequence logo generator. Genome Res 14: 1188–1190

Daems S, Ceusters N, Valcke R, Ceusters J (2022) Effects of chilling on the photosynthetic performance of the CAM orchid Phalaenopsis. Frontiers in Plant Science 13

Degenkolbe T, Giavalisco P, Zuther E, Seiwert B, Hincha DK, Willmitzer L (2012) Differential remodeling of the lipidome during cold acclimation in natural accessions of Arabidopsis thaliana. Plant J 72: 972–982

Degraeve-Guilbault C, Bréhélin C, Haslam R, Sayanova O, Marie-Luce G, Jouhet J, Corellou F (2017) Glycerolipid Characterization and Nutrient Deprivation-Associated Changes in the Green Picoalga Ostreococcus tauri. Plant Physiology 173: 2060–2080

Degraeve-Guilbault C, Gomez RE, Lemoigne C, Pankansem N, Morin S, Tuphile K, Joubès J, Jouhet J, Gronnier J, Suzuki I, Coulon D, Domergue F, Corellou F (2020) Plastidic Δ6 Fatty-Acid Desaturases with Distinctive Substrate Specificity Regulate the Pool of C18-PUFAs in the Ancestral Picoalga Ostreococcus tauri. Plant Physiol 184: 82-96

Degraeve-Guilbault C, Pankasem N, Gueirrero M, Lemoigne C, Domergue F, Kotajima T, Suzuki I, Joubès J, Corellou F (2021) Temperature Acclimation of the Picoalga Ostreococcus tauri Triggers Early Fatty-Acid Variations and Involves a Plastidial ω3-Desaturase. Front Plant Sci 12: 639330

Derelle E, Ferraz C, Rombauts S, Rouze P, Worden AZ, Robbens S, Partensky F, Degroeve S, Echeynie S, Cooke R, Saeys Y, Wuyts J, Jabbari K, Bowler C, Panaud O, Piegu B, Ball SG, Ral JP, Bouget FY, Piganeau G, De Baets B, Picard A, Delseny M, Demaille J, Van de Peer Y, Moreau H (2006) Genome analysis of the smallest free-living eukaryote Ostreococcus tauri unveils many unique features. Proc Natl Acad Sci U S A 103: 11647–11652

Domergue F, Abbadi A, Zahringer U, Moreau H, Heinz E (2005) In vivo characterization of the first acyl-CoA Delta6-desaturase from a member of the plant kingdom, the microalga Ostreococcus tauri. Biochem J 389: 483–490

Doyle JA, Donoghue MJ (1993) Phylogenies and angiosperm diversification. Paleobiology 19: 141–167

Fabri JHTM, de Sá NP, Malavazi I, Del Poeta M (2020) The dynamics and role of sphingolipids in eukaryotic organisms upon thermal adaptation. Progress in lipid research 80: 101063–101063

Gietz RD, Schiestl RH (2007) High-efficiency yeast transformation using the LiAc/SS carrier DNA/PEG method. Nat Protoc 2: 31–34

Gömann J, Herrfurth C, Zienkiewicz A, Ischebeck T, Haslam TM, Hornung E, Feussner I (2021) Sphingolipid long-chain base hydroxylation influences plant growth and callose deposition in Physcomitrium patens. New Phytol 231: 297–314

Gostincar C, Turk M, Gunde-Cimerman N (2010) The evolution of fatty acid desaturases and cytochrome b5 in eukaryotes. J Membr Biol 233: 63–72

Gronnier J, Germain V, Gouguet P, Cacas JL, Mongrand S (2016) GIPC: Glycosyl Inositol Phospho Ceramides, the major sphingolipids on earth. Plant Signal Behav 11: e1152438

Haak D, Gable K, Beeler T, Dunn T (1997) Hydroxylation of Saccharomyces cerevisiae ceramides requires Sur2p and Scs7p. J Biol Chem 272: 29704–29710

Hannich JT, Umebayashi K, Riezman H (2011) Distribution and functions of sterols and sphingolipids. Cold Spring Harb Perspect Biol 3

Haslam TM, Feussner I (2022) Diversity in sphingolipid metabolism across land plants. Journal of Experimental Botany 73: 2785–2798

Henderson GP, Gan L, Jensen GJ (2007) 3-D ultrastructure of O. tauri: electron cryotomography of an entire eukaryotic cell. PLoS ONE 2: e749

Hsu FF, Turk J, Zhang K, Beverley SM (2007) Characterization of inositol phosphorylceramides from Leishmania major by tandem mass spectrometry with electrospray ionization. J Am Soc Mass Spectrom 18: 1591–1604

Huby E, Napier JA, Baillieul F, Michaelson LV, Dhondt-Cordelier S (2020) Sphingolipids: towards an integrated view of metabolism during the plant stress response. New Phytol 225: 659–670

Igersheim A, Endress PK (1998) Gynoecium diversity and systematics of the paleoherbs. Botanical Journal of the Linnean Society 127: 289–370

Imai H, Ohnishi M, Hotsubo K, Kojima M, Ito S (1997) Sphingoid base composition of cerebrosides from plant leaves. Bioscience, biotechnology, and biochemistry 61: 351–353

Ishikawa T, Imai H, Maki KY (2014) Development of an LC-MS/MS method for the analysis of free sphingoid bases using 4-fluoro-7-nitrobenzofurazan (NBD-F). Lipids 49: 295–304

Ishikawa T, Ito Y, Kawai-Yamada M (2016) Molecular characterization and targeted quantitative profiling of the sphingolipidome in rice. Plant J 88: 681–693

Katoh K, Standley DM (2013) MAFFT Multiple Sequence Alignment Software Version 7: Improvements in Performance and Usability. Molecular Biology and Evolution 30: 772–780

Kawahara K, Moll H, Knirel YA, Seydel U, Zähringer U (2000) Structural analysis of two glycosphingolipids from the lipopolysaccharide-lacking bacterium Sphingomonas capsulata. Eur J Biochem 267: 1837–1846

Kay H, Grünewald E, Feord HK, Gil S, Peak-Chew SY, Stangherlin A, O’Neill JS, van Ooijen G (2021) Deep-coverage spatiotemporal proteome of the picoeukaryote Ostreococcus tauri reveals differential effects of environmental and endogenous 24-hour rhythms. Communications Biology 4: 1147

Lemoine F, Correia D, Lefort V, Doppelt-Azeroual O, Mareuil F, Cohen-Boulakia S, Gascuel O (2019) NGPhylogeny.fr: new generation phylogenetic services for non-specialists. Nucleic Acids Research 47: W260–W265

Li SF, Zhang GJ, Zhang XJ, Yuan JH, Deng CL, Hu ZM, Gao WJ (2016) Genes encoding Delta(8)- sphingolipid desaturase from various plants: identification, biochemical functions, and evolution. J Plant Res 129: 979–987

Li Y, Lou Y, Mu T, Ke A, Ran Z, Xu J, Chen J, Zhou C, Yan X, Xu Q, Tan Y (2017) Sphingolipids in marine microalgae: Development and application of a mass spectrometric method for global structural characterization of ceramides and glycosphingolipids in three major phyla. Anal Chim Acta 986: 82–94

Lingwood D, Simons K (2010) Lipid rafts as a membrane-organizing principle. Science 327: 46–50

Liu N-J, Hou L-P, Bao J-J, Wang L-J, Chen X-Y (2021) Sphingolipid metabolism, transport, and functions in plants: Recent progress and future perspectives. Plant Communications 2: 100214

López Alonso D, García-Maroto F, Rodríguez-Ruiz J, Garrido JA, Vilches MA (2003) Evolution of the membrane-bound fatty acid desaturases. Biochemical Systematics and Ecology 31: 1111–1124

Lou Y, Schwender J, Shanklin J (2014) FAD2 and FAD3 desaturases form heterodimers that facilitate metabolic channeling in vivo. J Biol Chem 289: 17996–18007

Mamode Cassim A, Grison M, Ito Y, Simon-Plas F, Mongrand S, Boutté Y (2020) Sphingolipids in plants: a guidebook on their function in membrane architecture, cellular processes, and environmental or developmental responses. FEBS Lett 594: 3719–3738

Marin B, Melkonian M (2010) Molecular phylogeny and classification of the Mamiellophyceae class. nov. (Chlorophyta) based on sequence comparisons of the nuclear- and plastid-encoded rRNA operons. Protist 161: 304–336

Mashima R, Okuyama T, Ohira M (2019) Biosynthesis of long chain base in sphingolipids in animals, plants and fungi. Future Sci OA 6: Fso434

Monnier A, Liverani S, Bouvet R, Jesson B, Smith JQ, Mosser J, Corellou F, Bouget FY (2010) Orchestrated transcription of biological processes in the marine picoeukaryote Ostreococcus exposed to light/dark cycles. BMC Genomics 11: 192

Moulager M, Corellou F, Vergé V, Escande ML, Bouget FY (2010) Integration of Light Signals by the Retinoblastoma Pathway in the Control of S phase Entry in the Picophytoplanktonic Cell Ostreococcus PLoS Genet

Nagano M, Ishikawa T, Ogawa Y, Iwabuchi M, Nakasone A, Shimamoto K, Uchimiya H, Kawai-Yamada M (2014) Arabidopsis Bax inhibitor-1 promotes sphingolipid synthesis during cold stress by interacting with ceramide-modifying enzymes. Planta 240: 77–89

Okino N, Li M, Qu Q, Nakagawa T, Hayashi Y, Matsumoto M, Ishibashi Y, Ito M (2020) Two bacterial glycosphingolipid synthases responsible for the synthesis of glucuronosylceramide and α- galactosylceramide. J Biol Chem 295: 10709–10725

Rennie EA, Ebert B, Miles GP, Cahoon RE, Christiansen KM, Stonebloom S, Khatab H, Twell D, Petzold CJ, Adams PD, Dupree P, Heazlewood JL, Cahoon EB, Scheller HV (2014) Identification of a sphingolipid α-glucuronosyltransferase that is essential for pollen function in Arabidopsis. Plant Cell 26: 3314–3325

Riva-Roveda L, Escale B, Giauffret C, Périlleux C (2016) Maize plants can enter a standby mode to cope with chilling stress. BMC Plant Biology 16: 212

Ronquist F, Teslenko M, van der Mark P, Ayres DL, Darling A, Höhna S, Larget B, Liu L, Suchard MA, Huelsenbeck JP (2012) MrBayes 3.2: Efficient Bayesian Phylogenetic Inference and Model Choice Across a Large Model Space. Systematic Biology 61: 539–542

Schlattmann P, Dirnagl U (2010) Statistics in experimental cerebrovascular research: comparison of more than two groups with a continuous outcome variable. J Cereb Blood Flow Metab 30: 1558–1563

Shah KH, Almaghrabi B, Bohlmann H (2013) Comparison of Expression Vectors for Transient Expression of Recombinant Proteins in Plants. Plant molecular biology reporter 31: 1529–1538

Shiels K, Tsoupras A, Lordan R, Nasopoulou C, Zabetakis I, Murray P, Saha SK (2021) Bioactive Lipids of Marine Microalga Chlorococcum sp. SABC 012504 with Anti-Inflammatory and Anti- Thrombotic Activities. Mar Drugs 19

Sperling P, Heinz E (2003) Plant sphingolipids: structural diversity, biosynthesis, first genes and functions. Biochimica et Biophysica Acta (BBA)-Molecular and Cell Biology of Lipids 1632: 1–15

Sperling P, Ternes P, Zank TK, Heinz E (2003) The evolution of desaturases. Prostaglandins Leukot Essent Fatty Acids 68: 73–95

Sperling P, Zahringer U, Heinz E (1998) A sphingolipid desaturase from higher plants. Identification of a new cytochrome b5 fusion protein. J Biol Chem 273: 28590–28596

Stonik VA, Stonik IV (2018) Sterol and Sphingoid Glycoconjugates from Microalgae. Mar Drugs 16

Tardif M, Atteia A, Specht M, Cogne G, Rolland N, Brugiere S, Hippler M, Ferro M, Bruley C, Peltier G, Vallon O, Cournac L (2012) PredAlgo: a new subcellular localization prediction tool dedicated to green algae. Mol Biol Evol 29: 3625-3639

Tavares S, Grotkjaer T, Obsen T, Haslam RP, Napier JA, Gunnarsson N (2011) Metabolic engineering of Saccharomyces cerevisiae for production of Eicosapentaenoic Acid, using a novel {Delta}5- Desaturase from Paramecium tetraurelia. Appl Environ Microbiol 77: 1854–1861

Ternes P, Wobbe T, Schwarz M, Albrecht S, Feussner K, Riezman I, Cregg JM, Heinz E, Riezman H, Feussner I, Warnecke D (2011) Two pathways of sphingolipid biosynthesis are separated in the yeast Pichia pastoris. J Biol Chem 286: 11401–11414

Tonon T, Sayanova O, Michaelson LV, Qing R, Harvey D, Larson TR, Li Y, Napier JA, Graham IA (2005) Fatty acid desaturases from the microalga Thalassiosira pseudonana. FEBS J 272: 3401–3412

Ukawa T, Banno F, Ishikawa T, Kasahara K, Nishina Y, Inoue R, Tsujii K, Yamaguchi M, Takahashi T, Fukao Y, Kawai-Yamada M, Nagano M (2022) Sphingolipids with 2-hydroxy fatty acids aid in plasma membrane nanodomain organization and oxidative burst. Plant Physiol 189: 839–857

Vaezi R, Napier JA, Sayanova O (2013) Identification and functional characterization of genes encoding omega-3 polyunsaturated Fatty Acid biosynthetic activities from unicellular microalgae. Mar Drugs 11: 5116–5129

Vítová M, Čížková M, Náhlík V, Řezanka T (2022) Changes in glycosyl inositol phosphoceramides during the cell cycle of the red alga Galdieria sulphuraria. Phytochemistry 194: 113025

Wang J, Chen YL, Li YK, Chen DK, He JF, Yao N (2021) Functions of Sphingolipids in Pathogenesis During Host-Pathogen Interactions. Front Microbiol 12: 701041

Yamashita S, Miyazawa T, Higuchi O, Takekoshi H, Miyazawa T, Kinoshita M (2022) Characterization of Glycolipids in the Strain Chlorella pyrenoidosa. J Nutr Sci Vitaminol (Tokyo) 68: 353–357

Zhou Y, Zeng L, Fu X, Mei X, Cheng S, Liao Y, Deng R, Xu X, Jiang Y, Duan X, Baldermann S, Yang Z (2016) The sphingolipid biosynthetic enzyme Sphingolipid delta8 desaturase is important for chilling resistance of tomato. Sci Rep 6: 38742

